# Differential proliferation regulates multi-tissue morphogenesis during embryonic axial extension: Integrating viscous modeling and experimental approaches

**DOI:** 10.1101/2024.02.26.581143

**Authors:** Michèle Romanos, Tasha Salisbury, Samuel Stephan, Rusty Lansford, Pierre Degond, Ariane Trescases, Bertrand Bénazéraf

## Abstract

The study of how mechanical interactions and different cellular behaviors affect tissues and embryo shaping has been and remains an important challenge in biology. Axial extension is a morphogenetic process that results in the acquisition of the elongated shape of the vertebrate embryonic body. Several adjacent tissues are involved in the process, including the tissues that form the spinal cord and musculoskeletal system: the neural tube and the paraxial mesoderm, respectively. Although we have a growing understanding of how each of these tissues elongates, we still need to fully understand the morphogenetic consequences of their growth and mechanical interactions. In this study, we develop a 2D multi-tissue continuum-based mathematical model to simulate and study how differential growth, tissue biophysical properties, and mechanical interactions affect the morphogenesis of the embryonic body during axial extension. Our model captures the long-term dynamics of embryonic posterior tissues previously observed *in vivo* by time-lapse imaging of bird embryos. It reveals the underestimated influence of differential tissue proliferation rates in inter-tissue interaction and shaping by capturing the relative impact of this process on tissue dynamics. We verified the predictions of our model in quail embryos by showing that decreasing the rate of cell proliferation in the paraxial mesoderm affects long-term tissue dynamics and shaping of both the paraxial mesoderm and the neighboring neural tube. Overall, our work provides a new theoretical platform to consider the long-term consequences of tissue differential growth and mechanical interactions on morphogenesis.

## Introduction

Differences in growth rate and biophysical properties have been proposed to be influential parameters for shaping biological forms since the last century (1). In vertebrate embryos, tissue morphogenesis occurs early during development when the body transforms from a round shape (disk or sphere, depending on the species) to a characteristic elongated one where the longest axis defines the anteroposterior direction (2,3). This extension involves the coordinated shaping of the different tissues that compose the embryonic body, such as the neural tube (NT)-which gives rise to the central nervous system, the paraxial mesoderm (PSM)-which gives rise to the vertebrae and muscles, and the notochord (NC)-an axial mesodermal tissue that secretes diffusible molecules involved in the patterning of both of these tissues to specify a variety of different cell types. Located in the embryo’s tail, posterior to these elongating tissues, lies an embryonic territory that contains an undifferentiated pool of progenitor cells called NMC for neuromesodermal competent cells (4–7). These cells self-renew, undergo specification, and migrate into the elongating neural tube and paraxial mesoderm, hence contributing to the extension of these tissues. Besides the addition of progenitor from the posterior zone (PZ), multiple cell processes are implicated in posterior tissue extension, for instance, cellular rearrangements such as convergent extension by intercalation have been shown to participate in the posterior extension of different posterior tissues, in different species (8,9). A gradient of non-directional motility within the paraxial mesoderm has also been shown to drive tissue extension in bird embryos. The transition of posterior high cell motility to low motility in the anterior part of the tissue is linked to a fluid-to-solid phase transition, which is proposed to play a key role in axial extension (10,11). Even though the relative importance of the action of each posterior tissue on the whole elongation process is still to be elucidated, experiments showing that posterior PSM deletion slows down elongation more than deletion of other tissues point out a pivotal role of this tissue in this process (10). Further studies recently proposed that paraxial tissue can generate forces to push on axial tissue, thus sustaining the extension process (12). Overall, the process of posterior extension is defined by a specific and complex choreography in which tissues grow differently and slide along one another while elongating (13). The neural tissue and the notochord, in particular, are moving toward the posterior part of the embryo faster than the paraxial mesoderm. Because axial extension occurs in growing embryos, the question of the role of cell proliferation and, more generally, differential growth involvement in this process arises. The answer differs significantly when looking at different species of vertebrate embryos. In zebrafish, the posterior extension does not rely on growth as it does not involve a massive change in tissue volumes and because cell cycle mutants do not display obvious truncation (14–16). On the contrary, in catfish, mice, and chick embryos, volumetric growth occurs in the developing tails (13,14). It has been shown that bird or mouse posterior embryonic tissues are actively proliferating with nearly all cells being engaged in the cell cycle possibly explaining most of this volumetric growth (17,18). However, measures of cell cycle durations in bird embryos have shown significant variations among posterior tissues (13). For instance, the average cell cycle duration is shorter in the paraxial mesoderm compared to the axial mesoderm or the neural tube. Although the role of cell proliferation on axial extension has been proposed to be minimal on a short-time duration in bird embryos (10), we still ignore whether it might influence multi-tissue morphogenesis on longer-time scales.

In the last two decades, several modeling approaches have been developed to enhance our comprehension of the underlying processes governing posterior axial extension. They successfully recapitulate and bring insight into many critical biological aspects of this complex morphological process. This includes the roles of FGF8 signaling on cell migration, posterior morphogenesis and patterning (19,10,20,21), the influence of cell adhesion on cell motion and tissue biophysical properties (22,23,11), as well as the putative mechanical coupling between posterior tissues (12). However, most of the current axial extension modeling approaches primarily center on the paraxial mesoderm’s role, neglecting a comprehensive exploration of how other developing tissues might influence the elongation process. Additionally, these approaches frequently disregard the significance of cell proliferation, placing greater emphasis on growth attributed primarily to cell injection from the progenitor zone.

This study aims to understand the role of differential growth and tissue interactions in coordinating the extension of the posterior tissues of the vertebrate embryo. In particular, we want to identify the most critical cell and tissue properties implicated in this coordination. Our approach to understanding multi-tissue growth dynamics takes a distinctly fully mechanical point of view, excluding assumptions about external signals and responses. To do so, we developed a new theoretical framework of continuum models that allowed us to test the influence of proliferation, injection, and viscosity in the different physically interacting posterior tissues of the embryo and to analyze their impact on elongation and tissue characteristics. We adopt a partial-differential-equation (PDE)-based model in a 2D multi-tissue framework, which integrates differential biophysical properties that contribute to the dynamics of the tissues. Our model builds upon previous works (24) by considering that tissue growth is driven by cell proliferation and cell ingression from the progenitor zone.

The main outcome of this study is that contrary to what is currently admitted in the literature, cell proliferation, rather than cell injection from the progenitor zone, is the most critical parameter in coordinating axial extension and shaping of posterior tissues. Our novel modeling approach for multi-tissue development not only highlights the role of differential tissue growth but also emphasizes the reciprocal mechanical influence that growing interacting tissues have on each other’s shape and elongation dynamics.

## Results

### Establishment of the continuum-based modeling and quantification of tissue cell entry from the progenitor zone

We aimed to develop a model describing the spatio-temporal dynamics of the PSM and neural tube densities. In the model, variations in cell densities are driven by tissue-specific cell proliferation rates (8.75 hrs for the PSM and 10.83 hrs for the NT) and cell ingression from the PZ as described in the literature. The PZ is modeled as a regressing injection zone along the anteroposterior axis, located at the posterior tip of the modeled tissues (Fig. 1A, B, Supplementary Materials and Methods). It participates in adding new cells at the posterior tip of both the NT and the PSM. Recent data (11–13) indicate that modeling posterior tissues as independent growing tissues, i.e., without mutual interactions, is not sufficient to explain how these tissues expand over time and suggest that intra-tissue and inter-tissue mechanics have consequences on tissue shape and dynamics. Therefore, we used a PDE-based model to account for the mechanical effects within and between neighboring tissues by considering different tissue velocities, which are strongly linked to tissue viscosity (Fig.1C). In this model, tissue-specific cell proliferation rates, cell injection, tissue densities, and viscosities are acting in concert to influence pressure and tissues’ densities, and therefore, tissue deformations. We set up the model with a portion of the NT already formed, flanked by two stripes of PSMs on either side. These tissues are constrained laterally by the lateral plate and anteriorly by the somites (two very dense structures), constituting the model’s fixed limits (walls). To make our model as close as possible to the dynamics of the vertebrate embryo, we collected parameters from the literature such as cell density, tissue viscosity, inter-tissue friction, and cell proliferation rates (Fig. 1D). For instance, the main parameter that was missing at the beginning of this study was the injection rate of new cells from the PZ feeding the growing tissues. To estimate this parameter, we analyzed the time-lapse movies of transgenic quail embryos (4) to measure how many cells per hour enter the PSM and the NT. We measured the average cell motion vectors within specific zones delimiting the interface between the PZ and the NT, as well as the PZ and the PSM in our model. We applied these motion vectors to the tissue density measured in these zones to estimate the flux of cells across these two inter-tissue boundaries (Fig. 1E, Supplemental Fig. 1, Supplemental Fig. 2). Our data show that, for a representative coronal tissue section of 10 um of thickness, 10 cells/hour on average enter the PSMs from the PZ whereas 5 cells/hour enter the neural tube. With the different parameters we collected and measured, we developed a biologically based theoretical framework that allows for modeling growth and forms of posterior tissues during bird embryonic development.

**Figure 1.**
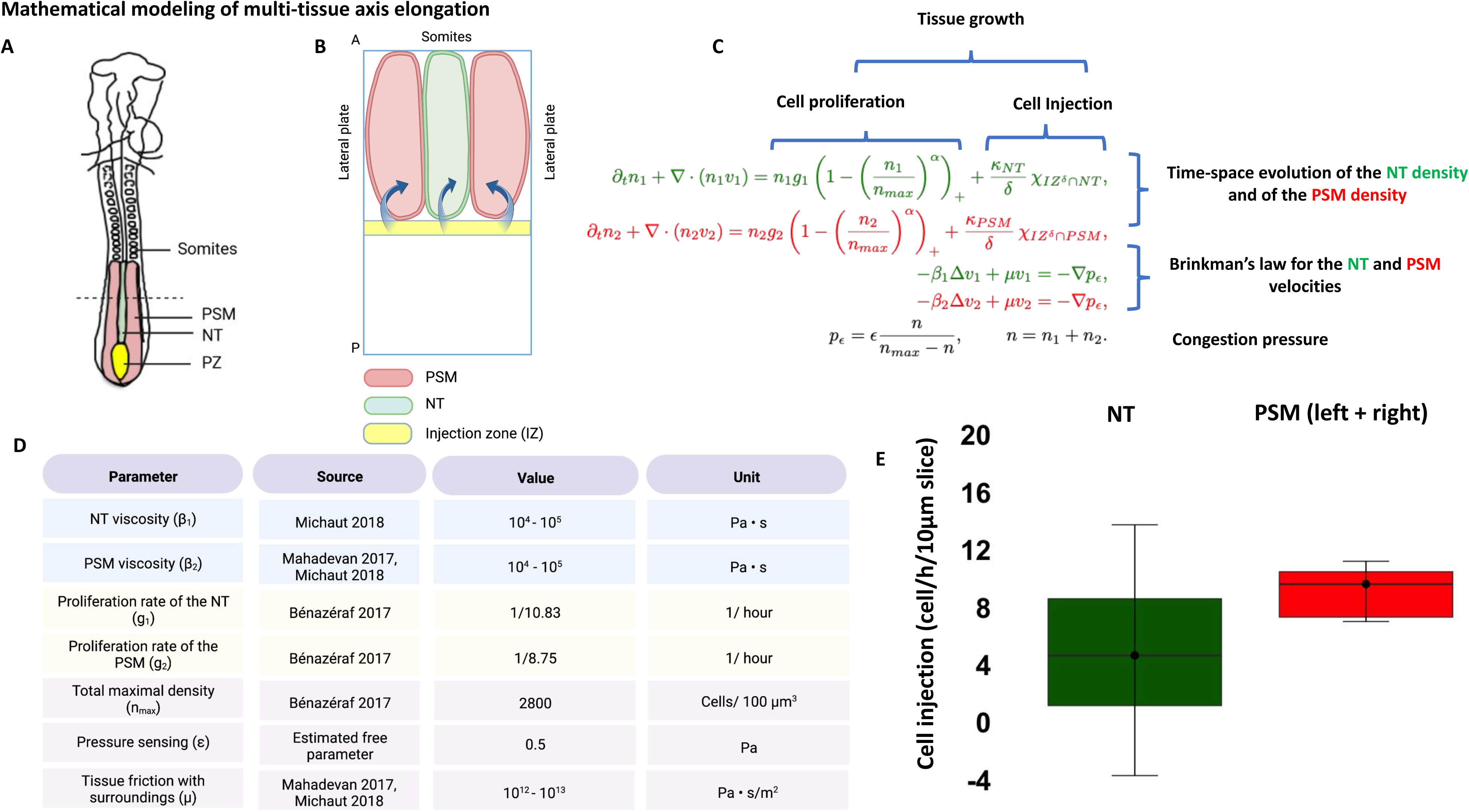
**A.** Schema of the vertebrate embryo. Posterior tissues are color-coded in red (PSM), green (NT), and yellow (PZ). **B.** Representation of the model setting. The NT (green) is flanked with two PSMs (red). Cell injection from the PZ (blue arrows) occurs at the posterior tip of the tissues (yellow zone). The lateral walls represent the lateral plate, and the upper wall (anterior) represents the last formed somite. **C.** The model equations. Equations in green refer to the dynamics of the NT and in red that of the PSM. The expression for the congestion pressure (in black) couples the NT and PSM equations through the Brinkman law for the velocities. **D.** List of model parameters, their source, value, and unit. **E.** Quantification of cell injection in the NT (green) and the left and right PSMs (red); y-axis represents the number of cells per hour entering the tissue from the PZ measured on a slice of depth 10 μm.

### Continuum-based modeling recapitulates key aspects of *in vivo* tissue dynamics

We first verified that our model can reproduce living embryos’ tissue growth and elongation features. We observed that after 20 hours, the model reproduces the tissue shapes comparable to the ones observed *in vivo* (Fig. 2A-B, Supplementary Movie 1). Tissues form and elongate posteriorly, and the cell density distributions show increasing gradients from posterior to anterior, as described in the embryo (10,13). We measured the elongation rate and found it equal to 128 um/hour (dashed line Fig 2B), which is in the range described in the literature (10,20). We plotted the velocity profiles within each tissue to investigate if the model recapitulates the global tissue dynamics observed *in vivo*. As in the embryo, we observed that the PSM and the NT display an anteroposterior velocity-increasing gradient (Fig. 2C, Supplementary Fig. 5B). Interestingly, the vectors located in the most posterior part of the tissue display medial-to-lateral biases which support the appearance of vortices in cell motions as it has been observed *in vivo* by quantification of rotational tissue movements clockwise and counterclockwise on the right and left PSM, respectively (13). The analysis of these rotational movements validates the presence of clockwise and counterclockwise movements in the PSM in our model (Fig. 2D). Furthermore, we noticed that within the anterior part of NT, the velocity vectors located close to the interface with the PSM point inwards, indicating a response to the PSM compression (Fig. 2C). To estimate potential pressure in the model we computed the pressure profile. We found out that the model displays pressure in the PSM that is globally higher than that in the NT, with a significant increase close to the interface with the NT (Fig. 2E, blue arrow, Supplementary Fig. 5A), and with an anteroposterior decreasing pressure gradient in both tissues (Fig. 2E). To check if this compression between tissues could be related to differential local tissue expansion, we analyzed the velocity divergence as a readout of expansion in our tissues (Fig. 3F). Interestingly, we found a posterior-to-anterior decreasing gradient of divergence in the PSM (green arrow) with a divergence that is globally higher than in the NT (blue arrows), as described *in vivo*. Finally, we wanted to test if our model can reproduce the inter-tissue sliding quantified using transgenic quail embryo imaging data (13). To do so, we quantified the differential motion between the PSM and the NT (i.e., sliding) (Fig. 2G). We defined and computed the sliding as the difference in the average NT and PSM velocities along the anteroposterior axis. Our analysis shows a significant (positive) sliding in the mid-to-anterior part of the tissues, reaching a maximum value of 12 um/h, with this sliding slowly decreasing posteriorly. The velocity of the NT is then much higher than that of the PSM, which causes the NT to slide past the PSM, as is the case *in vivo*. Overall, our results show that our continuum model successfully captures multiple aspects of the tissue dynamics, including rotational movements, expansion/compression profiles, and tissue sliding, all mirroring the *in vivo* processes, indicating that mesoscale growth and mechanics are an important part of the multi-tissue morphogenesis taking place in bird embryos.

**Figure 2.**
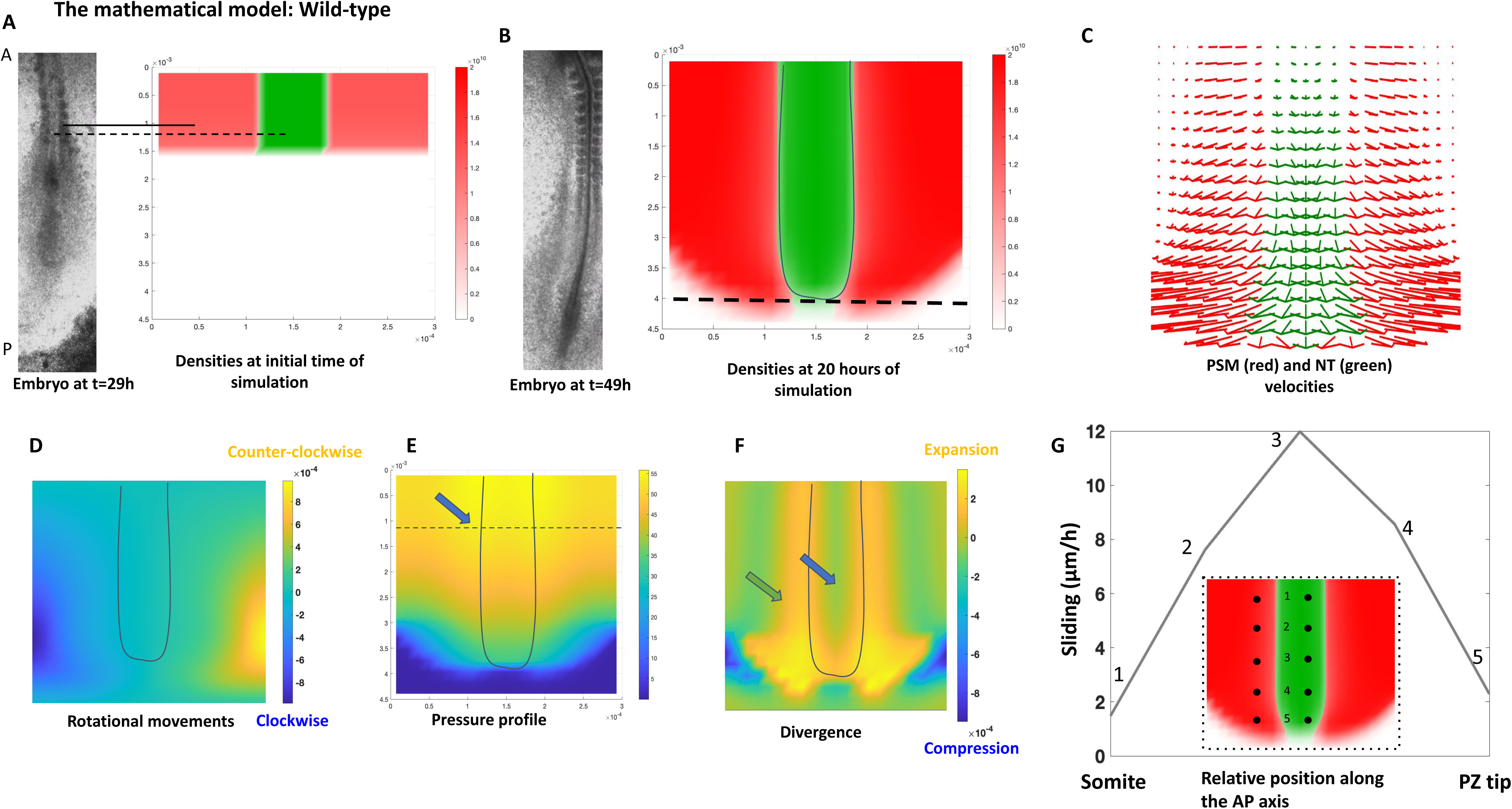
**A.** Left: quail embryo development at t=29 hours of development (see supplementary movie). Right: initial NT density (middle tissue) and PSM density (left and right) in the wild-type simulation of the mathematical model. The solid line shows PSM in the embryo, and in the simulation, the dashed line shows the NT. Notations A and P, respectively, denote the anterior and the posterior. **B.** Embryo at t=49 hours of development (left) and density plots in the wild-type simulation of the mathematical model after 20 hours of development. Colorbar represents density value (in IS units: cells/m^2). Solid line on density plot outlines the NT shape. Dashed line represents the elongation of the axis. **C.** Velocity vectors of PSM (red) and NT (green) in the simulation at t=20 hours. Vectors are plotted only in the areas that each tissue occupies. **D.** Rotational movements in the PSM: plot of Curl (v₂) at t=20 hours. Solid line on density plot outlines the NT shape. **E.** Pressure profile (p_ε) at t=20 hours. Colorbar gives pressure value in Pascal. The solid line on the density plot outlines the NT shape. The dashed line is a mediolateral cross section where we plot the pressure along this section in Supplementary Figure 5A. **F.** Tissue expansion and compression. Plot of Divergence (v₂) with representative colorbar. The solid line on the density plot outlines the NT shape. **G.** Sliding dynamics between NT and PSM computed over the last hour of simulation at 5 spots equally spaced along the antero-posterior axis defined by the axis somite-PZ tip. Inlet : density plot in (B) with the 5 spots (denoted 1 to 5) where sliding and tissue velocities are computed.

**Figure 3.**
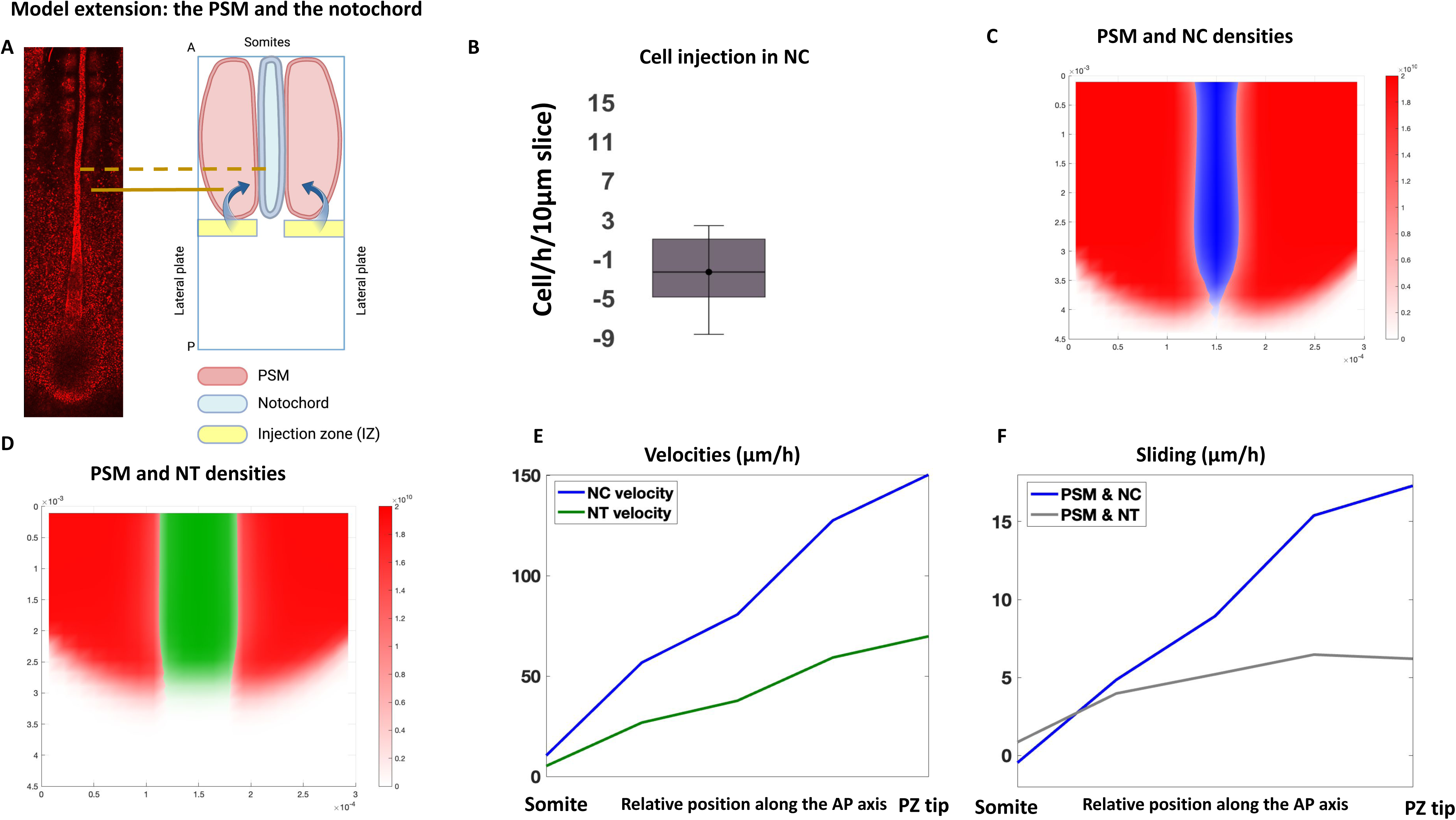
**A.** Left: transgenic quail embryo at t=29 hours of development with cell nuclei marked in red. Right: representation of the model setting. The Notochord (blue) is flanked with two PSMs (red). Cell injection from the PZ (blue arrows) occurs only at the posterior tip of the PSM (yellow zone). The lateral walls represent the lateral plate, and the upper wall (anterior) represents the last formed somite. Solid line shows PSM in the embryo and in the simulation, dashed line shows the Notochord. Notations A and P, respectively, denote the anterior and the posterior. **B.** Quantification of cell injection in the Notochord; y-axis represents the number of cells per hour entering the tissue from the PZ measured on a slice of depth 10 μm. **C.** PSM and Notochord densities at t=13 hours of development in the simulation, and **D.** Plot of PSM and NT densities in the wild-type simulation at t=13 hours for comparison. Color bars represent density values in SI units (cell/m^2). **E.** Notochord (NC) velocity (blue) computed over 5 spots equally spaced along the antero-posterior axis (somite-PZ tip) over the last hour of simulation, and NT velocity (green) for comparison. **F.** Sliding dynamics between the PSM and the Notochord (blue) over the 5 spots along the antero-posterior axis and the last hour of simulation, and sliding dynamics between PSM and NT (grey) for comparison.

### Versatility of continuum-based tissue modeling for axial extension

Since our model effectively recapitulates the NT and PSM dynamics, we wanted to test its ability to capture the dynamics between other posterior tissues. More specifically, we sought to model the growth dynamics of the notochord (NC) as, similarly to the NT, this tissue is known to physically interact with the paraxial mesoderm, yet it exhibits distinct growth and dynamic properties (Fig. 3A) (13). Indeed, the NC has an average cell cycle duration of 28 hours, which is much slower than other posterior tissues (8.75 hrs for the PSM and 10.83 hrs for the NT). Despite this discrepancy, it displays the highest posterior velocity during axial elongation, making it the tissue that slides the most with the PSM (13). We wanted to test if our model could explain this apparent paradox. Similar to the PSM and the NT, we needed to estimate the flux of new cells from the PZ into the NC to evaluate NC growth fully. Using the same image analysis method, we did not observe any influx of cells coming from the PZ to the NC, indicating that, at the analyzed stages, cells of the progenitor zone do not contribute, or only minimally to the NC growth (Fig. 3B). Therefore, we set the cell flux equal to zero for the NC in our model. The NC viscosity was estimated to be close to that of the NT as both tissues are epithelial tubes, and cell proliferation in the NC was set to 28 hours. We simulated the axial elongation of the PSM and the NC and deduced a NC elongation rate of 150 um/hr. The numerical simulation shows that the notochord maintains its elongated shape with a subtle anterior thinning, consistent with biological observations. Notably, the simulated shape of the notochord is thinner than that of the neural tube under the same conditions (Fig. 3C, D). When comparing the axial tissues in these conditions (NT vs NC), the plot of the velocity field reveals an anteroposterior increasing gradient in both the NC and the NT, with the NC velocity being remarkably higher than that of the NT (Fig. 3E). As a consequence, our model shows a significant sliding along the anteroposterior axis between the PSM and the NC, with the NC reaching much higher velocities and sliding past the PSM (Fig. 3F). In contrast to the PSM/NT sliding, the PSM/NC sliding is significantly greater, mirroring the observed differences observed *in vivo* for these tissues. Our results show that our model can reproduce the NC dynamics by adapting the parameters specifically for the NC.

While the growth of NC (cell proliferation and progenitor flux) is limited compared with PSM and NT, its elongation rate is relatively fast. Our model underscores the role of the interplay between growth dynamics and mechanical effects in driving the elongation of the NC. Paradoxically, the limited growth of the NC facilitates the conversion of pressure from its surrounding tissues into posterior movement. Indeed, whether we increase NC proliferation or decrease it in the PSM in our model, both scenarios result in a slower extension of the NC (data not shown). Overall, our model demonstrates its versatility by successfully capturing various dynamic properties observed *in vivo* across multiple posterior tissues.

### Modeling the effects of differential growth *vs* injection

One key question in the vertebrate embryo axial extension field is the relative influence of cell injection versus tissue-specific cell proliferation. To understand the effect and the role of cell proliferation versus cell injection, we built and compared two scenarios (numerical simulations): in the first scenario, we completely inhibited cell proliferation in the PSM, by setting the corresponding parameter to zero; in the second one, we completely inhibited the entry of new cells from the PZ into the PSM. We compared these simulations and looked at specific properties of tissue dynamics (Fig. 4A). We analyzed the elongation rates in the two scenarios and found that inhibiting cell proliferation resulted in a severely reduced elongation rate (reaching 64 um/h), compared with inhibiting cell injection, which yielded an elongation rate of 128 um/h which was comparable to the control case (Fig. 4A, B). As the global shapes of the tissues exhibit significant differences between the two scenarios, we quantified the tissue shape and density at the final time. By measuring the width of the NT in each scenario, we observed that in the absence of cell proliferation in the PSM, the NT gains in width all along the anteroposterior axis at the expense of the PSM that appears thinner compared to the control or to the case without cell injection in the PSM (Fig. 4A, C). This observation is consistent with the fact that in the absence of cell proliferation, we noted little to no pressure in the PSM (see Supplementary Fig. 5C), which normally participates in applying latero-medial forces on the NT and contributes to its posterior extension rather than its widening. Next, we analyzed the effects of inhibiting either PSM cell proliferation or cell injection on the inter-tissue dynamics. Our analysis of the tissues’ posterior extension showed that when inhibiting cell proliferation in the PSM, both the PSM’s and the NT’s velocities strictly diminish along the anteroposterior axis compared with the control case (Supplemental Fig. 4A-4B). These results suggest that cell proliferation promotes the motion of both tissues and supports the PSM’s mechanical role in NT elongation.

**Figure 4.**
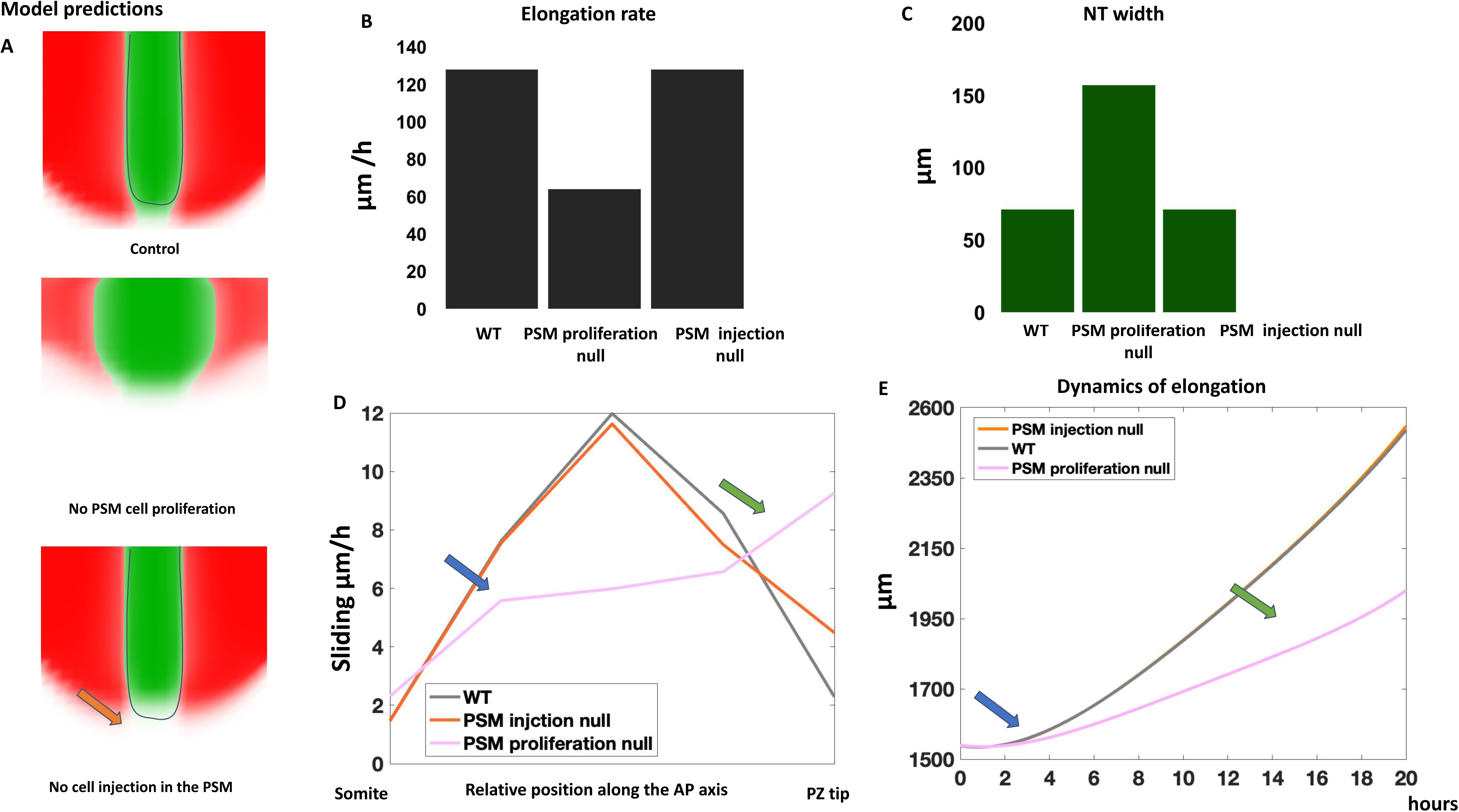
**A.** Upper panel: PSM and NT density plots at t=20 hours in the wild-type simulation, middle panel: PSM cell proliferation rate g_2 =0, lower panel: PSM injection rate κ_{PSM}=0. **B.** Elongation rate in the three simulations in (A). **C.** NT width for each simulation shown in (A). **D.** Sliding in each simulation computed over 5 spots equally spaced along the antero-posterior axis over the last hour of development in each simulation shown in (A). **E.** Fitting using a 4th order polynomial of PZ tip dynamics towards the posterior (IZ, see supplementary materials and methods) in each simulation shown in (A).

We next quantified the tissue sliding and noted that in the absence of cell proliferation (pink line), inters-tissue sliding is decreased anteriorly (Fig. 4D, blue arrow), and significantly increased at the most posterior positions along the anteroposterior axis (green arrow) in comparison with the control case (grey line). When we inhibited PSM cell injection, both PSM and NT velocities were slightly reduced but, more importantly, they were relatively similar to the control case along the anteroposterior axis (Supplemental Fig. 5C), which yielded a very low difference in sliding in comparison with the control case. However, the main difference is most noticed in the posterior position where the injection is taking place (Fig. 4D, orange vs gray line). Accordingly, we observed a difference in the posterior NT morphology, which was slightly wider in the no-cell injection case compared to the control (orange arrow Fig. 4A lowest panel). This highlights a very localized effect of cell injection on overall tissue morphology. Therefore, our model suggests that PSM cell proliferation orchestrates the dynamics and extension of both the NT and PSM, influencing tissue shaping and morphological changes along the anteroposterior axis, in contrast to the localized impact of cell injection.

To reconcile our data with the fact that inhibition of cell proliferation in the PSM on short-time ranges has been shown to have limited effects on extension (10,25), we explored the temporal dynamics of the influence of proliferation on PZ regression in our model. To monitor the axis elongation rate, we measured the evolution of the PZ posterior displacement over time (Fig. 4E, Supplementary Material and Methods). Strikingly, from 0 to 4.5 hours, the position of the PZ (blue arrow) was quite similar between the different considered cases (control, PSM cell proliferation null, PSM cell injection null). Over longer time scales, the differences in the PZ evolution became more significant, with the control having the most posterior position and the case with no PSM cell proliferation having the least posterior one (green arrow). Altogether, our results show the predominant growth effect of PSM cell proliferation compared to cell injection. Cell proliferation plays a vital role in maintaining tissue shape and density and promoting tissue velocity and posterior movements. It also affects inter-tissue dynamics, i.e., when absent, tissues experience less anterior sliding and, unexpectedly, more posterior sliding. Our analysis further reveals the strong coupling of the PSM and NT dynamics, i.e., when tissue-specific biological properties of one tissue are deregulated, it dramatically affects the other. Finally, our model suggests that the role of cell proliferation on tissue dynamics prominently takes hold with a certain latency as it starts manifesting clear effects only after at least 5-6 hrs.

### Differential cell proliferation plays a role in multi-tissue dynamics and tissue shaping

To validate the predictions of our model concerning the long-term effects of differential cell proliferation rates in multi-tissue morphogenesis, we experimentally inhibited cell proliferation in the PSM of transgenic quail embryos by electroporating a plasmid coding for the cell cycle inhibitor/CKI p27 in precursors of the paraxial mesoderm at stage HH5 embryo. Because with our electroporation technique, we obtain a transfection rate of 50% or higher in the PSM (7), inhibiting the cell cycle progression by artificially overexpressing p27 is expected to significantly slow down the average cell cycle duration of the whole tissue. To have access to global tissue dynamics, we performed time-lapse imaging in H2B-mcherry transgenic quail embryos electroporated with the p27 expression vector or control plasmids (Supplemental Movie 2). First, we verified that we effectively reduced the number of proliferating cells by executing an EdU staining on the electroporated embryos. Analysis shows that significantly fewer PSM cells electroporated with p27 have incorporated EdU after a 1-hour pulse compared to control cells (10% for p27 embryos vs 35% in control empty vectors) (Fig. 5A). Because we injected the p27-expression vector in progenitor cells before they enter the PSM we wanted to examine whether cell injection from the PZ was deregulated in the p27 electroporated embryos. To do so, we analyzed transgenic quail embryo movies by measuring the injection rate of p27 electroporated embryos vs. empty vector control embryos (described previously). We observed no significant difference in global cell injection in either the PSM or the NT (Fig. 5B). Therefore, our experimental condition corresponds to a situation in which proliferation is inhibited in the PSM tissue without significantly affecting the flux of cell injection from the PZ. We then analyzed tissue morphology and dimensions of embryos electroporated with the p27 and control vector. Analysis revealed that p27 electroporated embryos were shorter than control embryos; accordingly, PSM’s length was found reduced in the p27-electroporated embryos (Fig. 5C, D). Because our model predicts that inhibition of cell proliferation in the PSM affects NT morphology, we quantified the NT’s width at different locations along the AP-axis and found that the NT widens along the axis (Fig. 5C, E). We analyzed axis elongation and tissue motion in the movies of p27 or control vector electroporated transgenic quail embryos to study the tissue dynamics due to the inhibition of cell proliferation in the PSM. In line with the decrease in PSM size, we observed that elongation rates were decreased in the p27 condition compared with empty vector controls (45 vs 90 um/hrs). To assess whether tissue velocity is altered when PSM proliferation is diminished, we tracked cell motion in the NT and the PSM in the p27-electroporated embryo and compared it to control embryos along the AP-axis (Supplementary Fig. 3, Supplementary Fig. 4C-4D). We found that the PSM’s velocity is globally reduced along the anteroposterior axis in p27-electroporated embryos in comparison with the control embryos. We also found that the NT velocity of the p27-electroporated embryo is also globally reduced, especially anteriorly, and that posterior velocity is similar to that of the control embryos. By comparing the PSM and NT inter-tissue sliding, we found that, anteriorly, sliding is reduced in p27-electroporated embryos in comparison with the control ones (blue arrow). In contrast, posteriorly, the p27-electroporated embryos had a significantly higher sliding (green arrow) (Fig. 5G). Strikingly, our model accurately predicted the observed difference in sliding and tissue velocity behaviors in conditions where cell proliferation was inhibited in the PSM (Supplemental Fig. 3, Supplemental Fig. 4, compare Fig. 4D and 5G). In conclusion, our experimental approach shows that reducing the proliferation in the PSM leads to a decrease in the elongation of the PSM and the NT and also affects the shaping of the NT, highlighting the non-tissue autonomous mechanical action of the PSM on the NT. These results also validate the predictive power of our model to highlight the strong inter-tissue mechanical coupling during axis elongation.

**Figure 5.**
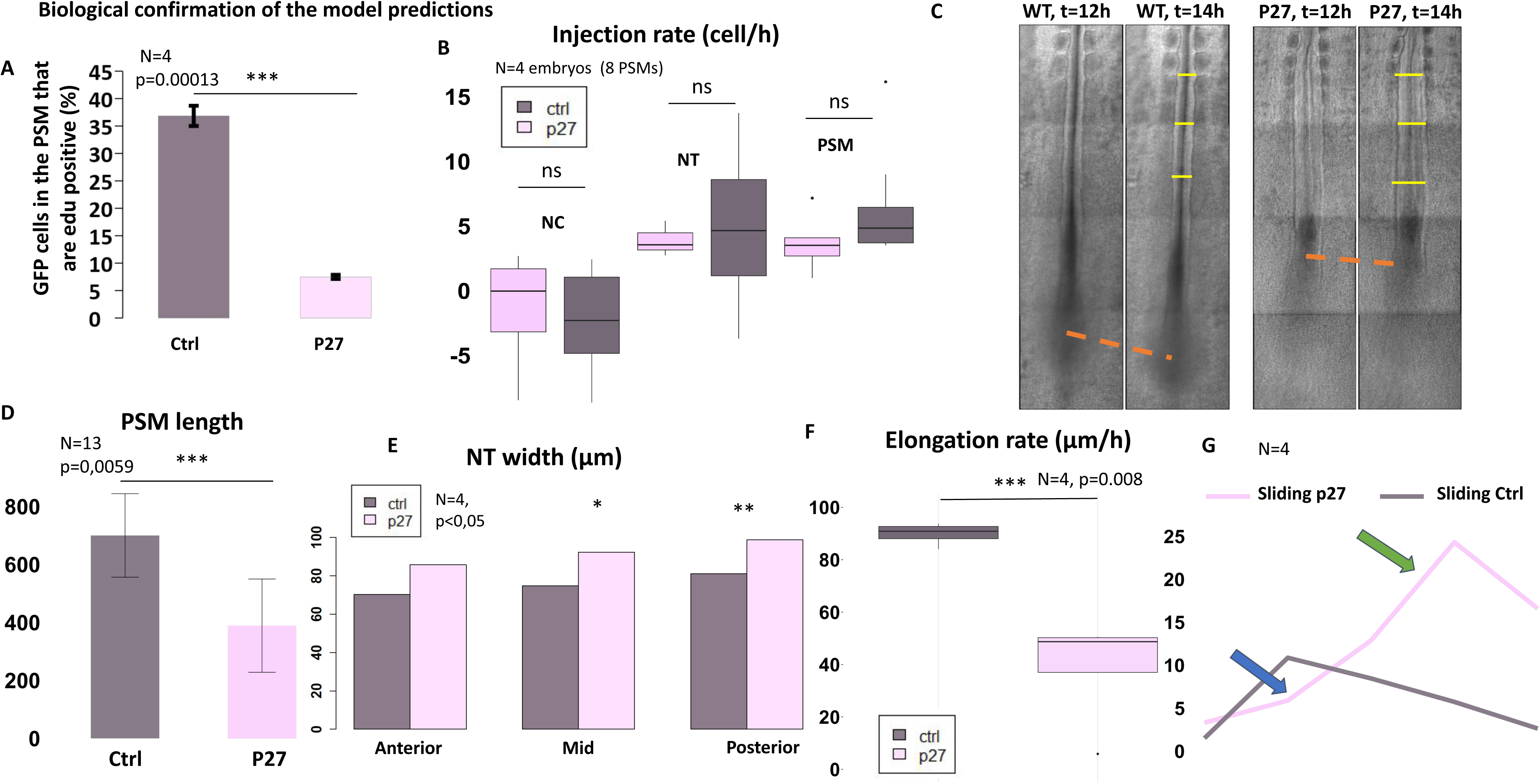
**A.** EdU positive cells in the PSM after electroporation of GFP (empty-grey) and GFP-p27 (pink). **B.** Quantifying injection rates in PSM, NT and Notochord in control and p27 embryos. ns = not statistical significant, see supplementary materials and methods for quantifications. **C.** Control (left panel) and p27 (right panel) quail embryos at t=29 hours (left) and t=49 hours (right) of development. Yellow bars along the NT represent tissue width. Orange dashed lines represent elongation. **D.** PSM length (measured in pixels) in control (grey) and p27 (pink) embryos. **E.** NT width (in μm) measured at three location on the antero-posterior axis: the anterior (at the last formed somite), middle (500 μm downwards for anterior position) and posterior (500 μm downwards from middle position) in control (grey) and p27 (pink) embryo at t=29 hours of development. (F) Elongation rate in control (grey) and p27 (pink) embryos measured over 2 hours of development. **G.** Sliding dynamics between NT and PSM velocities in control (grey) and p27 (pink) were computed over 5 spots equally spaced along the anteroposterior axis over the last hour of development. N=4; Statistical tests (t-tests) were performed, p-value and significance levels (*) shown.

## Discussion

In this manuscript, we developed and applied a new theoretical framework to study the influence of different types of growth and tissue mechanical properties on inter-tissue interactions and tissue shaping. We applied this model to axial extension, a complex multi-tissue morphogenetic process conserved between different vertebrate embryo species. The use of continuum modeling and hydrodynamic physics helped us characterize tissue dynamics and capture the essence of the underlying biological processes. One key feature of this new mathematical framework is that it allowed us to predict the relative influence of the different growth parameters on the outcome of the multi-tissue dynamics. In particular, we found that the proliferation rate was a prominent and underestimated feature of the long-term multi-tissue dynamics of axial extension, influencing both tissue extension and sliding between different tissues. Another interesting feature of this model is that it allows for the prediction and quantification of inter-tissue mechanical consequences and can efficiently predict how one tissue can shape its neighboring tissue. We indeed observed a mechanical impact of the PSM on the NT, suggesting that NT shape is not solely determined by inherent tissue-specific properties. Finally, we tested and validated the accuracy of our model’s prediction experimentally by demonstrating the implication of the higher proliferation rate of PSM cells compared to the ones of other tissues on multi-tissue dynamic shaping and axial extension.

To calibrate our model, we needed to characterize the injection rate from the progenitor region into the PSM, the NC, or the NT. Although the cellular contribution from the PZ to the PSM or the NT has been visualized/demonstrated by different fate mapping techniques at different developmental stages and in different species (6,7,26–28), it was still unclear how many cells entered the tissue by units of time. We have estimated the fluxes of cells entering the NT, PSM, and NC from the PZ in the quail embryo using cell/nucleï tracking to calibrate our model and enhance its fidelity to the vertebrate embryo. The analysis of our model indicates that the role of this injection alone is minimal compared to the role of cell proliferation and tissue mechanical interactions. Hence, the substantial contribution to elongation arises more from the proliferative activity of newly entered cells, generating significant offspring, rather than the injection itself. This is consistent with the fact that the effect of cell proliferation is observed on long-time scales and that the deletion of the progenitor region does not have immediate consequences on axis elongation speed compared with the deletion of the caudal PSM (10). However, the predominant role of cell proliferation in multi-tissue axis extension contrasts with previous data supporting the view that cell proliferation has little to no role in axis elongation (10, 25). This discrepancy might be explained by distinct experimental procedures used to inhibit cell proliferation so far. Inhibiting DNA synthesis, or cell division, which indeed affects proliferation, may only have a limited impact on cell growth in terms of volume gain. Indeed, our observation is that in aphidicolin-treated embryos, cells are larger than in control embryos, suggesting that tissue growth and the associated gain of volume might still be present in treated embryos (BB, unpublished observations). A second explanation could lie in the time scale of the analysis as the effect of inhibiting cell proliferation has been assessed on a rather shorter time scale (6 to 8 hours) in these previous works. Our model predicts that the effects of inhibiting cell proliferation become visible only after at least 8 hours of treatment exhibiting the time-dependent dynamics of cell and mechanical processes at the microscopic level and their manifestation at the macroscopic level. The delayed appearance of the phenotype could be attributed to the time required for changes in cell proliferation to manifest at the tissue density level and therefore influence intra-tissue and inter-tissue pressure. Interestingly, a recent study suggests that controlling cell density could be a crucial feedback mechanism in regulating elongation speed (29). It is important to note that the relative influence of cell proliferation in axis elongation may be dependent upon the developmental stage. Our analysis specifically concerns the interval encompassing stages 8HH to 12HH, corresponding approximately to the junctional neurulation process (30). It is plausible that in earlier developmental stages, particularly during primary neurulation, wherein gastrulation events are taking place, the influence of progenitor fluxes may have a more important impact on axial elongation than at later stages.

Since the PSM has been proposed to be a major driver of axis elongation, several agent-based models have been proposed to explain PSM elongation by itself (10,12,20). As we scale up to the tissue-level, the computational expense of agent-based models increases, giving way for partial differential equation (PDE)-based models. Although PDE models have proven useful in unraveling morphogenic principles in embryonic development (19–21,24,31,32), their emphasis remains on PSM elongation. The involvement of mechanical forces in axis elongation highlights the need to address both intra- and inter-tissue mechanical interactions. In our model, which focuses on elongation with expanding tissues, we performed a preliminary parameter sensitivity analysis to evaluate the importance of variations in biophysical properties relative to differences in proliferation (Supplemental Fig. 5E). The analysis results reveal that PSM and NT viscosity stand out as the second most critical parameters (after proliferation) in influencing the morphogenetic process of elongation. Aligned with this analysis, recent research has highlighted the critical role of tissue viscosity, particularly the graded viscosity within the PSM of the zebrafish embryo, in the fluid-to-solid transition along the antero-posterior axis, which determines the shape of the tissue and its ability to elongate unidirectionally (11,33). It is worth noting that our model system diverges from zebrafish, where tissue growth is restricted or absent during posterior body development, potentially explaining the disparities between the two models (14). Our continuum model, incorporating short and long-distance effects within and between highly proliferative tissues, reproduces embryonic tissue dynamics, including cell vortices resulting from our choice for the velocity viscous equation (24). Moreover, distinguishing tissues with differential growth potentials allowed us to predict the influence of the interactions between them, such as sliding or mutual shaping. Our analysis suggests that a growing tissue encountering the resistance of a neighboring growing tissue is crucial for correct shaping and elongation, with reciprocal effects from this growing tissue’s pressure. Therefore, our model emphasizes the importance of the geometric arrangement of interacting tissues in inter-tissue force transmission for axis elongation. Namely, the two PSMs must surround the NT to catalyze and channel its elongation, and as a consequence, the growing NT itself imposes forces back onto the PSMs. Future development of our multi-tissue model in 3 dimensions could further expand our understanding of the need for this particular geometrical arrangement. Moreover, a 3D model could allow exploring the simultaneous mechanical relationships between the different embryonic tissues: PSM/NT/NC. The fact that tissues slide relative to each other points out the importance of the nature of the physical interface between tissues. Several works have emphasized the role of the extracellular matrix, which accumulates at the boundaries between tissues, as an important player in the process of axis elongation. For instance, it has been shown that in mouse or zebrafish embryos cell/ECM interaction is involved in posterior tissue shaping and elongation (23,34,35). In particular, morpholino injection for integrin alpha 5 and alpha V in zebrafish leads to a bent NC, suggesting that if the link between developing tissue is impaired, one tissue can deform the other in a non-physiological manner (23). This study suggests that the interface between differential growing posterior tissues has to allow them to slide one against the other. It is interesting to note that in other developmental contexts such as gut morphogenesis, and because tissues are physically attached, they can deform, bend, or coil in a physiological manner instead of sliding. In that case, the tube and the mesentery, being attached and having different growing rates, promote the gut to coil (36).

This study presents a novel multi-tissue modeling framework in the context of morphogenesis. While our investigation primarily focuses on the vertebrate embryo, the polyvalence of this model extends its applications to various biological systems. Despite potential variations in specific mechanisms reflected by different parameter values, the model’s inherent generality makes it adaptable to diverse contexts. Moreover, our model goes beyond developmental biology and exhibits potential applications in oncology, particularly in studying cancers with different properties (viscosity, growth…). Its ability to accommodate tissues with distinct features positions it as a powerful tool for exploring the dynamics of heterogeneous tumors (37,38). Our model serves as a benchmark in multi-tissue modeling within developmental biology. Its minimal assumptions on tissue dynamics provide a unique approach to comprehending large-scale dynamics and morphologies. This distinctive feature highlights the model’s versatility as a robust framework for investigating a broad spectrum of developmental processes and extends its relevance to the broader field of systems in biology.

## Material and methods

### Quail embryo and embryo culture

The USC aviary, and local French hatchery (les cailles de Chanteloup) provided wild-type quail embryos (Coturnix japonica). PGK1:H2B-chFP quail lines have been described previously (39) and are maintained in the USC aviary. The embryos were staged according to (40) and Hamburger and Hamilton (41). Embryos were cultured *ex ovo* with filter paper on albumen agar plates using the EC (early chick) technique (42).

### Plasmids, electroporation, and proliferation essays

Mouse P27 construct was acquired through Addgene (plasmid 15192). We collected stages HH5-7 quail embryos. Empty or p27 vectors, (2–5 μg/μl) were microinjected between the vitelline membrane and the epiblast (43) in the anterior part of the primitive streak and surrounding epiblast in a region that contains paraxial mesoderm precursors (44). The electrodes were positioned on either side of the embryo, and five pulses of 5.2 V, with a duration of 50 ms, were carried out at a time interval of 200 ms. The embryos were screened for fluorescence and morphology. Proliferation rates were assessed by Edu staining (Click-iT EdU Alexa Fluor 647 Imaging Kit, Thermo Fisher Scientific, <u>C10340</u>)

### Imaging and image analysis

For live-imaging experiments, embryos were cultured at 37°C in culture imaging chambers (Lab Tek chambered #1 coverglass slide, Thermo Fisher Scientific) pre-coated with a mix of albumen agar (42). Embryos were imaged using an inverted 780 Zeiss microscope using confocal excitation with 20×/0.8, or 25×/0.8 objectives. For time-lapse imaging, several adjacent XYZ fields of view were stitched together post-collection. Images were taken every 5-10 µm in the z-axis with a time resolution ranging from 5 to 6 min. The Spot and track functions of Imaris were used to localize nuclei in the 3D image data set and to extract nuclei coordinates (x,y,z,t) (details on sliding quantification and referential used can be found in the supplementary methods). The data were then analyzed using Excel, MATLAB, or Python.

### Mathematical modeling

The mathematical model was simulated using a finite volume scheme on a staggered grid. All the numerical simulations were performed using Matlab 2022a. For more information refer to :

## Acknowledgements

The Benazeraf team members, particularly Cathy Soula and Myriam Roussigné, are gratefully acknowledged for their assistance and comments on the work. Financial support for this project was provided by the CNRS through the 80|Prime program (S.S/B.B), Ph.D. Fellowship funded by Toulouse University the Occitanie Region and the FRM (4th year) (M. Romanos). Special thanks are extended to David Huss for his valuable assistance in transgenic quail husbandry.

---

**Supplementary Figure 1.**
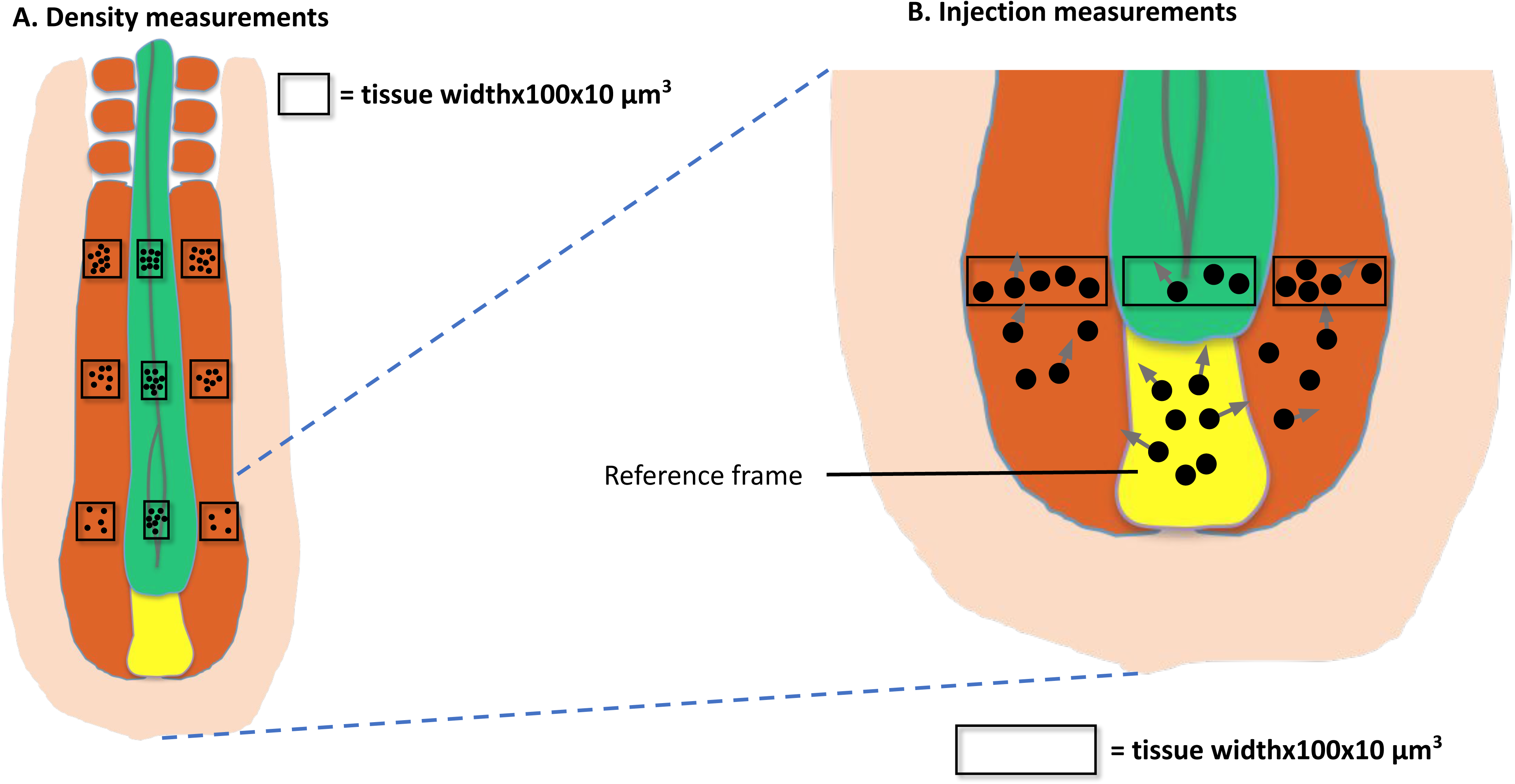
**A.** Schema of the posterior tissues: PSM in red, NT in green, PZ in yellow and lateral plate in light pink. Black squares represent the three regions considered to measure cell density (cell count). Three regions were considered in each tissue: anterior, mid and posterior to evaluate cell density along the antero-posterior axis. Size of region in each tissue: width = tissue width, length = 100 μm, depth = 10 μm. Black dots represents cells counted in each region (see supplementary materials and methods for quantifications). **B.** Zoom on the schema of posterior tissues. Regions considered for cell injection quantification have the same size as the ones for density measurements. Regions were placed at the most posterior part of each tissue (see supplementary materials and methods for quantifications).

**Supplementary Figure 2.**
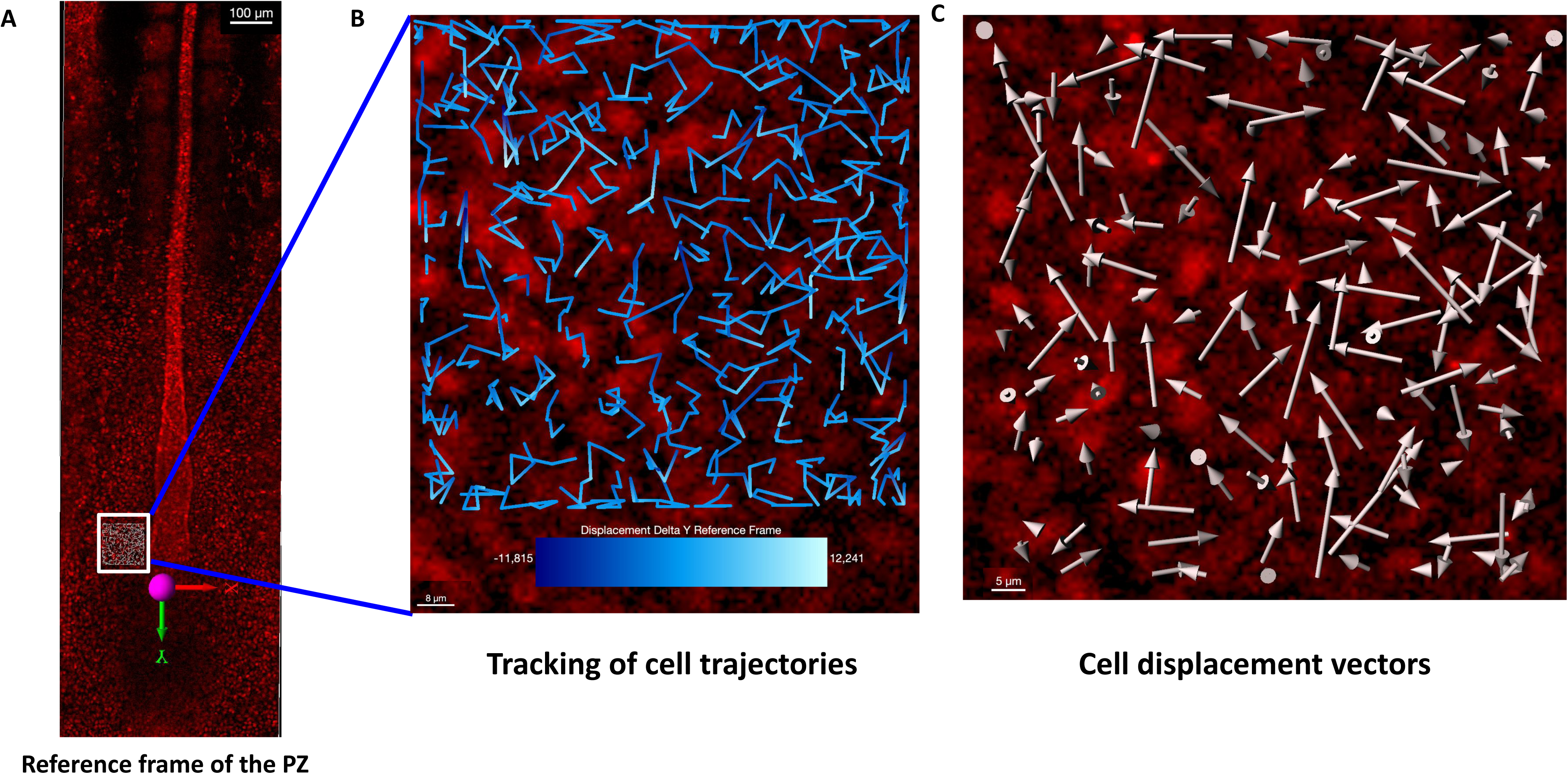
**A.** Screenshot of transgenic quail embryos (on Imaris). Cell injection was computed in the white square region taken at the posterior tip of the PZ (see supplementary materials and methods for quantifications) in the reference frame of the PZ (represented on the PZ). **B.** Tracking of cell trajectories (dragon tail on Imaris). Colorbar represents position on the AP axis with respect to the reference frame. Blue portions of trajectories represent cells exiting the PZ towards the anterior. **C.** Cell displacement vectors pointing towards the anterior representing cell trajectories from the posterior towards the anterior.

**Supplementary Figure 3.**
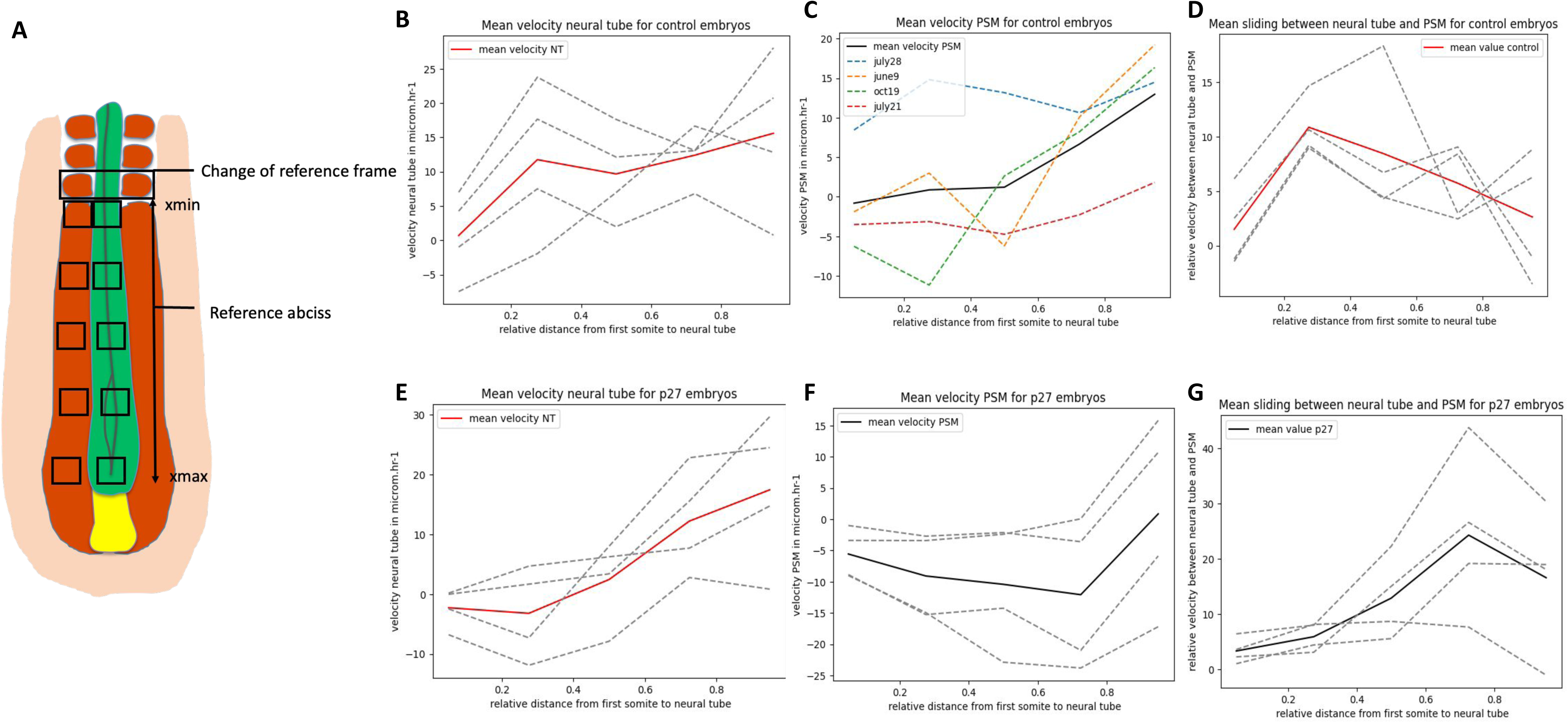
**A.** Schema of the posterior tissues. Quantifications of tissue velocities was done by changing the reference frame to that of the last formed somite. The position of the last formed somite is denoted x_min and the position of the NT tip is denoted x_max. The velocities are computed on 5 points equally spaced along the reference axis [x_min, x_max] for each embryo (see supplementary materials and methods). **B.** NT velocities in the WT embryos for N=4 embryos computed on the reference axis (dashed grey lines). The means of the velocities at each point on the axis are represented in the red solid line. **C.** PSM velocities in the WT embryos for N=4 embryos computed on the reference axis (dashed lines). The means of the velocities at each point on the axis are represented in the black solid line. **D.** Sliding between the NT and the PSM in the WT embryos: the difference between the NT and PSM velocities on each point of the reference axis for N=4 embryos are in dashed lines and the mean difference (sliding) is represented on the red solid line. **E.** NT velocities in the p27 embryos for N=4 embryos computed on the reference axis (dashed grey lines). The means of the velocities at each point on the axis are represented in the red line. **F.** PSM velocities in the p27 embryos for N=4 embryos computed on the reference axis (dashed lines). The means of the velocities at each point on the axis are represented in the black solid line. **G.** Sliding between the NT and the PSM in the p27 embryos: the difference between the NT and PSM velocities on each point of the reference axis for N=4 embryos are in dashed lines and the mean difference (sliding) is represented on the black solid line.

**Supplementary Figure 4.**
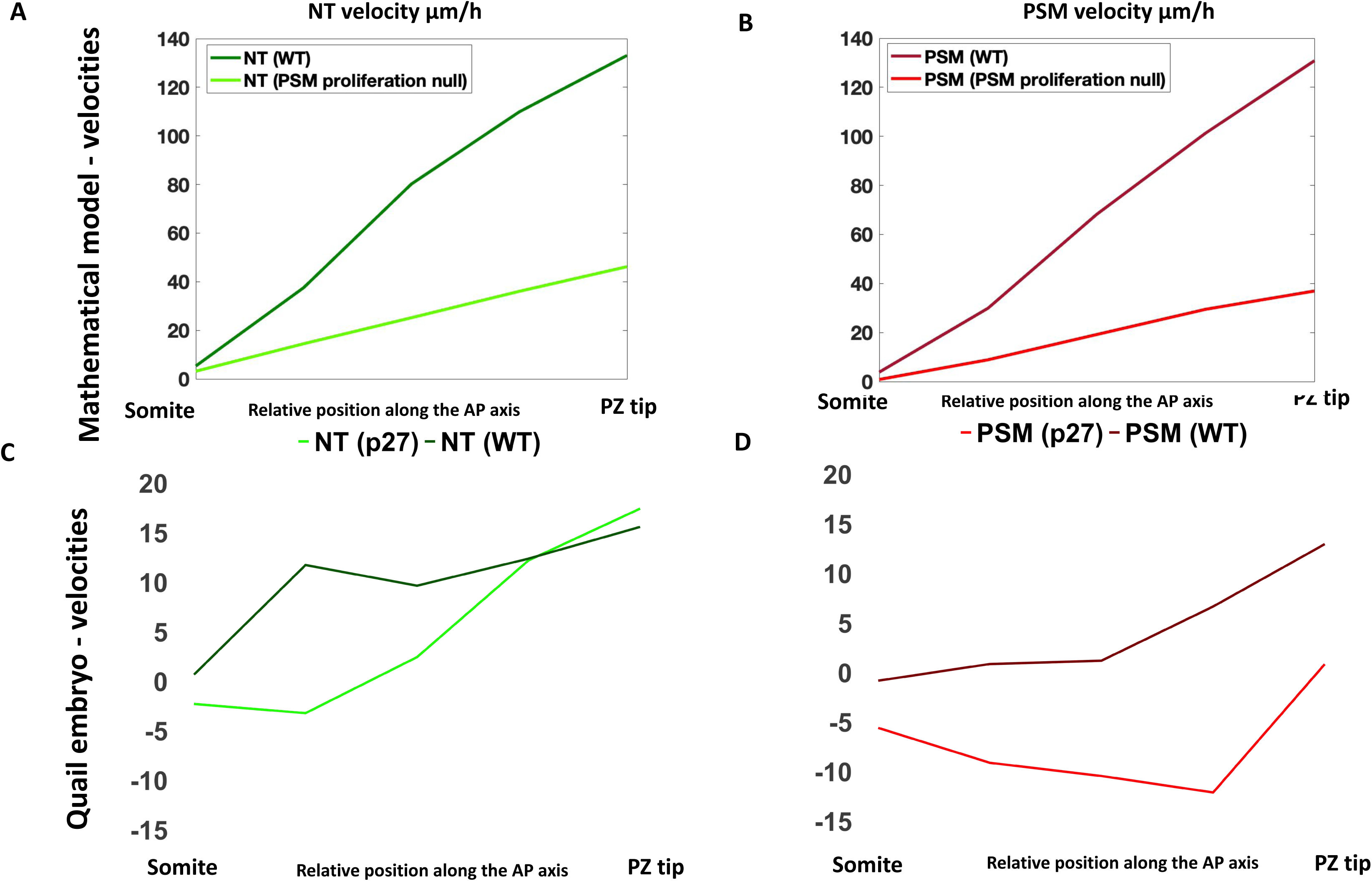
**A.** NT velocities in the WT simulation (dark green) and in the p27 simulation (light green). **B.** PSM velocities in the WT simulation (dark red) and in the p27 simulation (light red). **C.** NT velocities in the WT embryos (dark green) and in the p27 embryos (light green). **D.** PSM velocities in the WT embryos (dark red) and in the p27 embryos (light red).

**Supplementary Figure 5.**
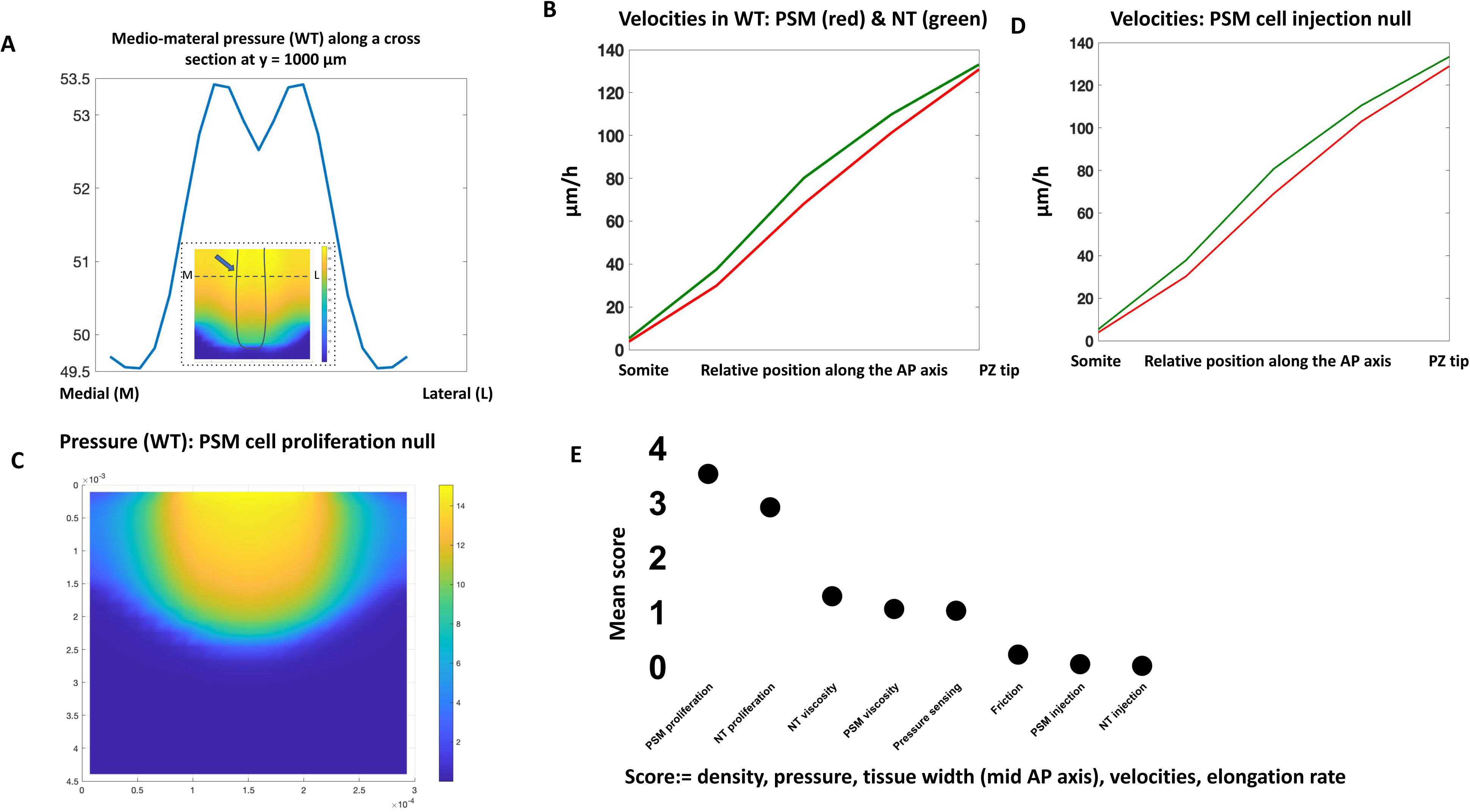
**A.** Pressure profile along a mediolateral cross section at y=1000 μm (on the AP axis) represented in the inlet (dashed line in the density plot). **B.** NT (green) and PSM (red) velocities in the WT simulation. **C.** Pressure profile in the simulation when cell proliferation in the PSM is null (g_2=0). **D.** NT (green) and PSM (red) velocities in the simulation when cell injection in the PSM is null (κ_{PSM} =0). **E.** Sensitivity analysis scoring: each parameter (shown on the x-axis) was deviated from its original (wild-type) value of ±10%, ±20%,±30%,±40%,±50%. Model outputs scored: PSM and NT densities, velocities, tissue width, pressure, and elongation rate. The mean score S(P) for each parameter (black dots) is calculated as the average score across all simulations corresponding to variations in that specific parameter (see supplementary material and methods).

## List of Supplementary movies

- Supplementary movie 1 : WT simulation
- Supplementary movie 2 : movie transgenic quail WT (left) and p27 (right)

Supplementary Movie 1: Simulation of the WT case. The NT (in green) is flanked with two PSMs (red). The title indicates the time in hours. The numerical domain is of length 4500 micrometers and of width 300 micrometers. The x and y axes are in meters. The tissues are color coded by their densities, the colorbar on the left indicates the density level (gradient from white to red): red = high density and white = no density. For clarity of the movie and figures only the colorbar for the PSM is represented (the NT colorbar is a gradient from white (no density) to green (high density)). Simulation parameters are detailed in the supplementary materials and methods and main Figure 1.

Supplementary Movie 2: Two movies of H2B-mcherry transgenic quail embryos Tg(PGK1:H2B-chFP) electroporated with the control plasmids (left) or p27 (cell cycle inhibitor/CKI p27) expression vector (right). Each cell nucleus is stained in red. EdU positive cells are stained in yellow. Scale in micrometers is represented on the lower left position of the movie, time in hours is represented in the lower right position.

