## Supplementary Materials and Methods for "Differential proliferation regulates multi-tissue morphogenesis during embryonic axial extension: Integrating viscous modeling and experimental approaches"

### Supplementary text

February 12, 2024

**1 Mathematical model** **1**

#### 1 Mathematical model

##### 1.1 Main variables and mathematical setup

We consider two cell population densities $n_1(t, x, y)$ (representing the NT) and $n_2(t, x, y)$ (representing the PSM) evolving in time $t$ and in space $(x, y)$. We opt for a 2D spatial model, describing a $10\mu m$-thick cross-section of the 3D embryonic tissues. The choice of a 2D spatial representation reflects that the depth of the tissues can be considered small compared with other lengths (average depth of the NT and PSM is $50\,\mu m$). It is also consistent with the fact that most of our biological data at hand are in 2D.

In addition to its density $n_i(t,x,y)$, each tissue is described through its local velocity vector $v_i(t,x,y)$, evolving in time and space. Furthermore, we will consider the total pressure $p(t,x,y)$ exerted in both tissues indistinctly. To sum up, our main variables are:

- $n_1(t,x,y)$: cell density of NT,
- $n_2(t,x,y)$: cell density of PSM,
- $v_1(t,x,y)$: tissue velocity of NT,
- $v_2(t,x,y)$: tissue velocity of PSM,
- $p(t,x,y)$: total pressure of the tissues.

These quantities are defined over a rectangular domain $\Theta = [0,l] \times [0,L]$, corresponding to a zone confined between the two lateral plates (at $x=0$ and $x=l$) and posteriorly to the last formed somites (at $y=0$).

#### 1.2 Equations for the dynamics of the tissues

The dynamics of the tissues is modeled by the following equations, for all $t \geq 0$ and $(x,y) \in \Theta$,

$$\partial_t n_1 + \nabla \cdot (n_1 v_1) = n_1 g_1 \left(1 - \left(\frac{n_1}{n_{max}}\right)^{\alpha}\right)_+ + \frac{\kappa_1}{\delta}\, \chi_{IZ^\delta \cap NT}, \tag{1}$$

$$\partial_t n_2 + \nabla \cdot (n_2 v_2) = n_2 g_2 \left(1 - \left(\frac{n_2}{n_{max}}\right)^{\alpha}\right)_+ + \frac{\kappa_2}{\delta}\, \chi_{IZ^\delta \cap PSM}, \tag{2}$$

$$-\beta_1 \Delta v_1 + \mu v_1 = -\nabla p, \tag{3}$$

$$-\beta_2 \Delta v_2 + \mu v_2 = -\nabla p, \tag{4}$$

$$p = \epsilon \frac{n}{n_{max} - n}, \qquad n = n_1 + n_2. \tag{5}$$

These equations include tissue mechanical properties (viscosity, friction, pressure), cell proliferation and injection from the PZ: the detailed modeling assumptions for each of them are given below.

**Mechanical properties.** We consider that both tissues are viscous, and confer them with viscosity parameters, respectively $\beta_1$ and $\beta_2$. In (3) and (4) we use the Brinkman law to govern the velocities $v_1$ and $v_2$ of each tissue. The parameter $\mu$ represents the friction between the tissue and its surroundings. The friction coefficients for the PSM and the NT are both taken equal to $\mu$: we made this assumption as, to our knowledge, the friction of the NT with its surrounding tissues is not a known parameter in literature and has not been measured.

In addition, the pressure law (5) is a function of the total density (the sum of $n_1$ and $n_2$), which allows tissue segregation (if segregation is imposed at initial time $t = 0$), [3]. The pressure is taken singular at $n = n_{max}$ which represents the maximal density inside the tissues measured *in vivo*. This form of the pressure law provides an upper bound on the cell densities, which is particularly relevant in the context of vertebrate embryo development and the tissues we want to model ([3], [2]).

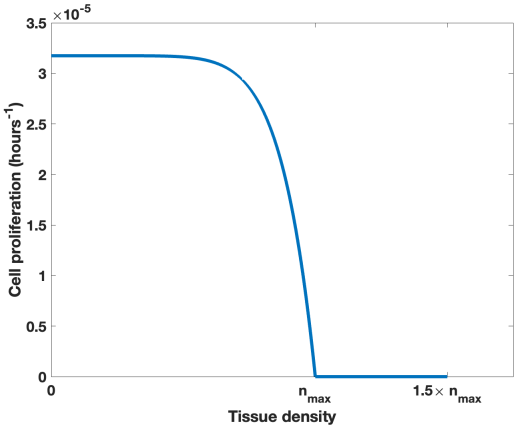

Figure 1: Plot of the cell proliferation function given by $g_2\left(1-\left(\frac{n_2}{n_{max}}\right)^{\alpha}\right)_+$ in Equation(2) with respect to the tissue density $n_2$ ranging between 0 and $1.5 \times n_{max}$.

**Cell proliferation.** In the evolution equations on the densities (1)-(2), the first term of the reaction terms (on the right-hand side) represents cell proliferation. The growth rates are taken as (reversed) sigmoidal functions with a saturating density equal to $n_{max}$. The steepness of the sigmoid is determined by the exponent $\alpha$. The parameter $g_1$ (respectively, $g_2$) represents the intrinsic proliferation rate of the NT (respectively, the PSM), that is, the theoretical proliferation rate in absence of saturating effects. The saturation effect accounts for the transition of cellular type, as well as possible dorso-ventral migration, in most anterior, most dense, regions of the NT and PSM where 3D structures such as the somites start to form.

**Cell injection.** Still in equations (1)-(2), the second term of the right-hand-side accounts for cell injection from the PZ. The flux of new cells from the PZ into each of the tissues is denoted $\kappa_1$ and $\kappa_2$ in respectively the NT in (1) and the PSM in (2). These new cells are added only at the elongating tip of each tissue in a region denoted $IZ^{\delta} \cap NT$ for the region of entry of new cells in the NT and $IZ^{\delta} \cap PSM$ for that of the PSM[1], see Figure 2. We define the entry zone as a thin rectangular of width $\delta$ that moves over time, translating according to the elongation dynamics. More precisely, as the PZ is attached to the NT at its posterior tip, we determine the posterior tip denoted $y_{tip}$ as the following,

$$y_{tip}(t) := \text{argmin}_{y\in[0;L]} \left\{ n_1(t, x_{mid}, y) \geq \frac{n_{max}}{2} \right\},$$

where $x_{mid} = l/2$ is the central position in the NT along the lateral axis. We then define the entry zones as,

$$IZ^{\delta}(t) \cap NT = [a_1; b_1] \times \left[ y_{tip}(t) - \frac{\delta}{2}; y_{tip}(t) + \frac{\delta}{2} \right],$$

with $[a_1; b_1]$ the width of the zone initially occupied by the NT, and,

$$IZ^{\delta}(t) \cap PSM = ([a_2; b_2] \cup [a_2'; b_2']) \times \left[ y_{tip}(t) - \frac{\delta}{2}; y_{tip}(t) + \frac{\delta}{2} \right],$$

[1]In equations (1)-(2) $\chi$ is the indicator function.

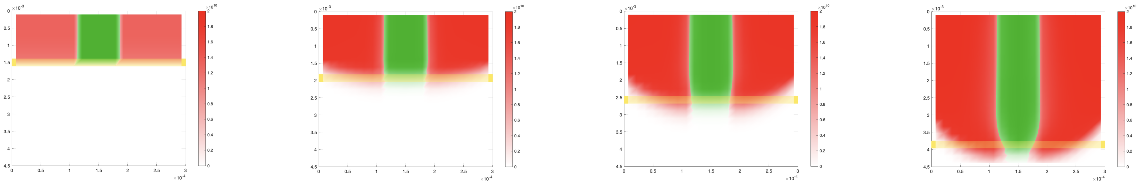

(a) $T = 0$ hours. (b) $T = 6.4$ hours. (c) $T = 12$ hours. (d) $T = 20$ hours.

Figure 2: Evolution of the injection zone (yellow rectangle) of the NT and PSM: $IZ^{\delta} \cap (NT \cup PSM)$ in the wild-type simulation.

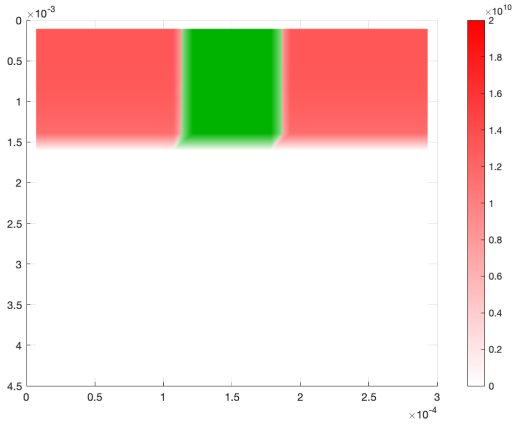

Figure 3: Plot of the densities at initial time: the NT (middle tissue) flanked with two PSM on the right and on the left.

with $[a_2; b_2]$ and $[a_2'; b_2']$ the width of the zone initially occupied by respectively the left and the right PSM (of equal width), see Figure 2. Values for $a_i$ and $b_i$, $i = 1, 2$ and for $a_2', b_2'$, as well as more details on the initial condition are given in Section 1.3.

**Remark 1** (Modeling of the PZ)**.** *The PZ could as well be modeled as a third tissue, leading to a model with three population densities: this alternative framework would introduce more modeling hypotheses, more parameters, and overall more complexity to our study. Here our strategy is to focus on the PSM and the NT and to not consider the PZ per se but only its contribution to the formation of those two tissues as an injection zone.*

#### 1.3 Numerical simulation

**Numerical setting and initial condition.** The numerical domain $\Theta$ is taken as the rectangle $[l \times L] = [300\mu m \times 4500\mu m]$. We consider the embryo with a portion of the NT and of the PSM already formed: we consider that the NT initially occupies a zone of length $L/3 = 1500\mu m$ and of width $71\mu m$. It is centered in $x$, that is, $[a_1; b_1] = [114\mu m; 185\mu m]$, and it is flanked with two PSM on the right and on the left, with length $L/3 = 1500\mu m$ and width of $100\mu m$ each, which corresponds to $[a_2; b_2] \cup [a_2'; b_2'] = [0\mu m; 100\mu m] \cup [200\mu m; 300\mu m]$. Experimental data indicate that the density in the NT is approximately constant and higher than that of the PSM which displays a clear antero-posterior density gradient [1]. We define $\overline{n_1}$ as the measured density in the

NT, $\overline{n_2}$ as the maximum density quantified in the PSM anteriorly and $\underline{n_2}$ as the minimal density quantified in the PSM posteriorly, see Section 3.1. As per our measurements of the tissue densities (see Section 3.1), and measurements in literature by [1], we have,

$$\overline{n_1} = \frac{0.017\, cells}{\mu m^2}, \qquad \overline{n_2} = \frac{0.013\, cells}{\mu m^2}, \qquad \underline{n_2} = \frac{0.011\, cells}{\mu m^2}.$$

We take the initial NT density as follows,

$$n_1(t=0,x,y) = \overline{n_1}\chi_{0\le y\le L/3}.$$

As for the PSM, we impose the density gradient on the initial condition, as follows,

$$n_2(t=0,x,y) = \overline{n_2}\chi_{0\le y\le L/8} + \left(\overline{n_2} - \gamma(y-L/8)^2\right)\chi_{L/8\le y\le L/3},$$

where $\gamma = \frac{\overline{n_2}-\underline{n_2}}{(L/3-L/8)^2}$ is computed such that the PSM density antero-posterior gradient ranges between $\underline{n_2}$ and $\overline{n_2}$.

The initial condition is shown in Figure 3.

**Remark 2.** *In the main and supplementary text we write all the quantities (e.g densities, length, width) in physically relevant units (cells/micrometer, micrometer). The units in the figures are in the international unit system (cells/meter, meter).*

**Boundary conditions.** The lateral numerical walls correspond to the lateral plate, while the upper wall, at the anterior of the vertebrate body, corresponds to the last formed somites. Considering all those as dense structures, we set the no flux boundary conditions for the densities, as well as homogeneous Dirichlet conditions for the velocities (no-slip condition) on all the boundaries, that is,

$$v_i = (v_{i_x}, v_{i_y}) = 0, \quad (n_i v_i)\cdot\nu = 0, \; i=1,2, \text{ on } \partial\Theta.$$

These conditions state that no density can flow out of the walls, and that when cells come in contact with these walls they strongly adhere to them. The posterior boundary is a theoretical wall, all simulations are stopped before the tissues reach this boundary.

**Model parameters.** We give here the values of the model's parameters used in the numerical simulations for the wild-type embryo. We first recall the parameters and the values (or ranges) that could be quantified experimentally. References for parameters found in the literature are provided, and for parameters we measured, the protocol measurements are detailed in the corresponding sections.

- $g_1 \approx \frac{1}{10.83}$ $[hour^{-1}]$. The cell proliferation rate in the NT was measured in [1] using EdU.
- $\kappa_1 \approx 0.046$ $[cells \cdot hour^{-1} \cdot \mu m^{-1}]$. Using image analysis on transgenic quail embryos, we measured on the Imaris software the entry of new cells from the PZ into the NT over a region of width equals to tissue width, length equal to $100\mu m$ and depth equals to $10\mu m$. The method is detailed in the corresponding section.
- $\beta_1 \approx 10^4 - 10^5\,[Pa \cdot s]$. The viscosity of the NT was measured in [5] (PhD thesis) using pipette aspiration technique. We note here that $\beta_1 > \beta_2$ according to [5], meaning that the NT is more viscous than the PSM.
- $g_2 \approx \frac{1}{8.75}$ $[hour^{-1}]$. The cell proliferation rate in the PSM was measured in [1] using EdU.

- $\kappa_2 \approx 0.128$ $[cells \cdot hour^{-1} \cdot \mu m^{-1}]$ (in the left and right PSM). Using image analysis on transgenic quail embryos, we measured on the Imaris software the entry of new cells from the PZ into the PSM over a region of width equals to tissue width, length equal to $100\mu m$ and depth equals to $10\mu m$. The method is detailed in the next section.

- $\beta_2 \approx 10^4 - 10^5$ $[Pa \cdot s]$. The viscosity of the PSM was measured in [7], [6], [5] using pipette aspiration technique.

- $\mu \approx 10^{12} - 10^{13}$ $[Pa \cdot s \cdot m^{-2}]$. The friction between the PSM and its surrounding tissues (NT, lateral plate, ectoderm, endoderm) was estimated in [7], [6], [5].

- $n_{max} \approx 0.02$ $[cells \cdot \mu m^{-2}]$. We measured the maximal densities in the tissues (see Section 3.1 for protocol). It can also be deduced from [1] where an averaged density map was represented along the antero-posterior axis of the vertebrate embryo, see [1].

- $l$ and $L$ are equal to $300\mu m$ and $4500\mu m$ respectively.

- $\delta$: we choose the width of the entry zone $\delta$ to be equal to one mesh cell length.

- $\epsilon = 1$. This free parameter controls the congestion pressure $p$. It was fixed such that we obtain the right elongation speed of the tissues.

- $\alpha = 8$. This parameter controls the steepness of the sigmoid growth function. It was fixed such that we obtain the right elongation speed of the tissues.

For the numerical simulations we take the following values:

| $g_1$ | $g_2$ | $\kappa_1$ | $\kappa_2$ | $\beta_1$ | $\beta_2$ | $\mu$ | $n_{max}$ | $l$ | $L$ | $\delta$ | $\epsilon$ | $\alpha$ |
|---|---|---|---|---|---|---|---|---|---|---|---|---|
| 1/10.83 | 1/8.75 | 0.046 | 0.128 | $3 \cdot 10^4$ | $2 \cdot 10^4$ | $10^{12}$ | 0.02 | 300 | 4500 | 214.2857 | 1 | 8 |

**Numerical scheme.** The mathematical model was simulated using a finite volume scheme on a staggered grid. All the numerical simulations were performed using Matlab 2022a. We use $N^2 = 21^2$ mesh cells, with space steps $\Delta x = \frac{l}{N} = 14.2857\mu m$ and $\Delta y = \frac{L}{N} = 214.2857\mu m$. The time step is dependent on the CFL (Courant-Friedrichs-Lewy) condition which varies at each iteration depending on the velocity, for details on the implementation and the numerical scheme see [3]. We simulate until we reach the final time set to $T = 20$ hours of developmental time. Simulation frames are saved every 6 minutes (of developmental time) to generate the supplementary movies, which corresponds to the time between frames in the time-lapse movies of quail embryos.

#### 1.4 The mathematical model for the PSM and the notochord

To simulate the PSM and the Notochord, the model equations (1)–(5) remain unchanged, with now $n_1$ and $v_1$ standing for the density and velocity vector of the Notochord. The only modifications are the values of the parameters, now adapted to the Notochord. This implies taking the injection rate (previously denoted as $\kappa_1$) in the Notochord equal to zero, as measured *in vivo* (main manuscript Figure 3). We change the proliferation rate (previously $g_1$) and take it equal to 1/23.5 hours, as measured and averaged in [1]. Finally, we adjusted the initial density in the Notochord, measured and detailed in Section 3.1, resulting in a value of $0.016\, cells/\mu m^2$. The viscosity parameter of the Notochord remains equal to that of the NT as they are both epithelial and tubular.

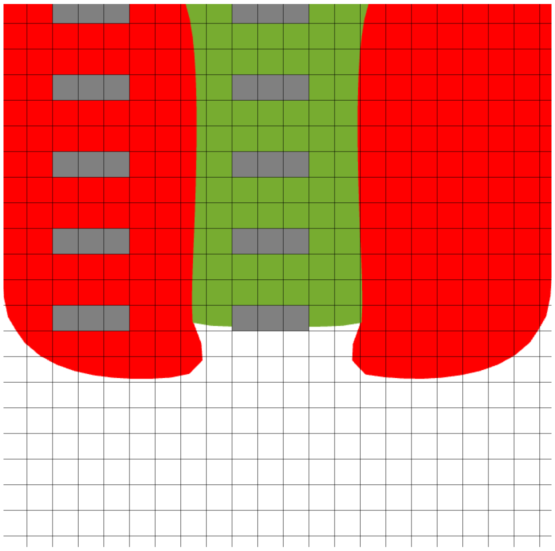

Figure 4: Schema of the tissues with the numerical mesh in the background. To obtain the central velocities, we compute the average velocities in the PSM and the in the NT on the mesh cells colored in gray respectively in mid PSM and mid NT and along the antero-posterior axis.

### 2 Quantification of model outputs

#### 2.1 Velocity, sliding and elongation quantification

To plot the tissue velocities (main manuscript Figures 3E,Supplementary Figure 4A-B, Supplementary Figure 5B-D) and the sliding (main manuscript Figures 2G, 3F, 4D) between the tissues (NT and PSM, NC and PSM) we computed the central velocities in each tissue, defined below. The method we use is very close to the analysis done in [1], which facilitates the comparison of numerical simulations and *in vivo* quantifications.

**Central velocity.** We compute the central velocity at five given heights $y_{j_0}, ...y_{j_4}$, equally spaced along the antero-posterior axis. For each given height $j_k$, we consider three mesh cells $(i_0-1, j_k)$, $(i_0, j_k)$ and $(i_0+1, j_k)$ centered within the width of the NT, see Figure 4. We then compute the mean central velocity of the NT at $y = y_{j_k}$ as,

$$V^1(y_j) = \frac{v^1_{i_0-1,j} n^1_{i_0-1,j} + v^1_{i_0,j} n^1_{i_0,j} + v^1_{i_0+1,j} n^1_{i_0+1,j}}{n^1_{i_0-1,j} + n^1_{i_0\,j} + n^1_{i_0+1,j}}, \text{ for } j = j_0, ..., j_4.$$

The plot of the velocity profile is generated based on the velocities averaged over the final hour of the simulation to accurately compare with the velocity quantification in the vertebrate embryo (see Section 3.3). We do similar computations for the PSM and Notochord velocities.

**Sliding.** To obtain the sliding in Figure 2G (main manuscript), we subtract the NT central velocity computed previously from the PSM central velocity, and we plot this difference. The sliding in Figure 3F (main manuscript) is obtained similarly with the NT central velocity replaced with the Notochord central velocity.

**Elongation dynamics.** To plot Figure 4E (main manuscript) which depicts the dynamics of elongation, we simply plot the evolution of $y_{tip}(t)$ defined in 1.2.

#### 2.2 Sensitivity analysis and scoring

A sensitivity analysis consists in varying individually each model parameter, while keeping all the others unchanged, and quantify the changes in the outputs thus obtained. We conducted a sensitivity analysis on the model by varying each of the parameters: $g_1$, $g_2$, $\beta_1$, $\beta_2$, $\epsilon$, $\mu$, $\kappa_1$, $\kappa_2$ by a reduction/increase of $10\%, 20\%, 30\%, 40\%, 50\%$ of their original values (wild-type) determined in Section 1.3. To each variation of a single parameter corresponds a simulation, and for each simulation, we evaluate the deviation of the model outputs from the wild-type simulation (simulated with the parameters from Section 1.3). For all simulations we assess the model outputs at the same time point $T_0 = 17$ hours, which corresponds to the end of the validity of the fastest simulation (before the tissues encounter the posterior boundary of the numerical domain). The quantified output changes are the following:

- The profiles of the NT and PSM densities ($n_1$ and $n_2$)
- The profile of the pressure $p$
- The profile of the PSM and NT lateral velocities $v_{1_x}$, $v_{2_x}$
- The profile of the PSM and NT posterior velocities $v_{1_y}$, $v_{2_y}$
- The NT and PSM widths
- The elongation rate

For each fixed parameter $\mathcal{P}$, we denote by $\mathcal{P}_0$ the value of the parameter used in the wild-type simulation (listed in Section 1.3) and by $\mathcal{P}_i$ the deviated values. More precisely,
$\mathcal{P}$ = parameter type
$\mathcal{P}_0$ = parameter value in wild-type simulation
$\mathcal{P}_i$ = value increased (or decreased) of $i \times 10\%$, for $i = -5 \dots 5$, that is, $\mathcal{P}_i = \mathcal{P}_0(1 + i/10)$.

For any given output $F$ among the outputs listed above, and any given parameter $\mathcal{P}$, we denote by $F(\mathcal{P}_0)$ the value of the output of the wild-type simulation and by $F(\mathcal{P}_i)$ the value of the output of the simulation with the deviated parameter $\mathcal{P}_i$. With these notations, we assign for each output $F$ the following score.

If $F$ is defined as a function on the whole domain $\Theta$ (velocities, densities, pressure), we take,

$$\text{Output score}(\mathcal{P} \mapsto F) = \sqrt{\frac{1}{10} \sum_{i \in \{-5,..,5\} \setminus \{0\}} \left( \frac{\mathcal{P}_0}{\|F(\mathcal{P}_0)\|_2} \frac{\|F(\mathcal{P}_0) - F(\mathcal{P}_i)\|_2}{|\mathcal{P}_i - \mathcal{P}_0|} \right)^2},$$

where $\| \cdot \|_2$ is the $L2$ norm. If $F$ is a scalar output (width, elongation rate) we take,

$$\text{Output score}(\mathcal{P} \mapsto F) = \sqrt{\frac{1}{10} \sum_{i \in \{-5,..,5\} \setminus \{0\}} \left( \frac{\mathcal{P}_0}{|F(\mathcal{P}_0)|} \frac{|F(\mathcal{P}_0) - F(\mathcal{P}_i)|}{|\mathcal{P}_i - \mathcal{P}_0|} \right)^2}.$$

Finally the parameter score $S(\mathcal{P})$ is based on the root mean square of all the output scores, more precisely,

$$S(\mathcal{P}) = \text{Parameter score}(\mathcal{P}) := \sqrt{\sum_{F \text{ output}} \left(\text{Output score}(\mathcal{P} \mapsto F)\right)^2}.$$

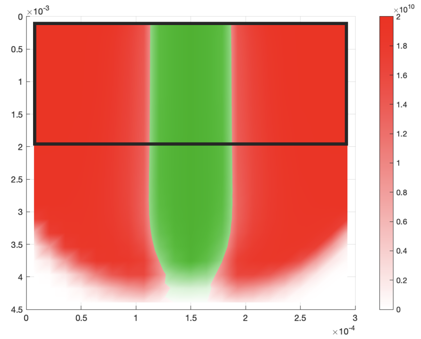

Figure 5: Schema of the tissues. The black contoured area represents the fixed section $sec_{WT} = [0, 1900] \times l$ in the Wild-type case used for the computation of the score.

This yields the score represented in Supplementary Figure 5E.

**Remark 3.** *When comparing simulations at $T_0 = 17$ hours, some exhibit a severe reduction in the elongation rate (e.g reducing the PSM growth by 50%), resulting in shorter tissues. To compare tissue densities, pressure, and velocities with those of the wild-type simulation, the comparison is restricted to regions occupied by the tissues. The aim is to overlap the two simulations (Wild-type and deviated) as closely as possible, at least over a limited area. This is achieved by computing the output scores $S(\mathcal{P})$ only on sections of fixed dimension ($1900\mu m \times l$) that vary along the antero-posterior axis. We fix the section in the wild-type case as $sec_{WT} = [0, 1900] \times l$ (Figure 5) and vary the section in deviated simulations along the axis to evaluate which one gives the best score for the deviated simulation. We then choose the best (minimal) score $S$ given by the section. This yields,*

$$S(\mathcal{P}) = \min_{a_i \in [0, L-1900]} \{S(\mathcal{P})_{|sec_{WT}, sec_{var}}\},$$

*where $S(\mathcal{P})_{|sec_{WT}, sec_{var}}$ is computed over $sec_{WT} = [0, 1900] \times l$ for the wild type and $sec_{var} = [a_i, a_i + 1900] \times l$ for the section in the deviated simulation.*

### 3 Quantifications in the vertebrate embryo

In what follows we provide the details of how density and cell injection measurements were acquired using Imaris software.

#### 3.1 Density measurements

We used quail embryos with DAPI staining to measure tissue densities in the PSM, NT and Notochord on the Imaris software (using the Spot function). In each tissue, we measure the total number of cells inside regions of length $100\mu m$, of width that of the tissue, and of depth $10\mu m$ in the PSM, NT and Notochord. These cross-sections were taken in the mid-depth of the tissue. Measurements were taken in three locations along the antero-posterior axis: anterior (at the level of the somite), mid ($800\mu m$ downwards from the somite) and posterior ($800\mu m$ downwards from the middle location).

#### 3.2 Injection measurements

We used live movies of transgenic quail embryos Tg(PGK1:H2B-chFP) where each cell nucleus is stained [4, 1], to quantify cell injection from the PZ into the PSM (main manuscript Figure 1E), the NT (main manuscript Figure 1E) and the Notochord (main manuscript Figure 3B) on the Imaris software (using Spot and Track functions). First, we changed the reference frame from the laboratory frame to the frame of the moving posterior tip to better capture cell exit. This can be achieved by averaging cell velocities inside the PZ and shifting the reference frame to follow that of the average velocity computed. We defined regions with a length of $100\mu m$, a width equal to that of the tissue (denoted as $w$), and a depth of $10\mu m$ within the PSM, NT, and Notochord. These cross-sections were taken in the mid-depth of the tissue. They were strategically positioned at the most posterior part of each tissue to effectively capture cell entry. We tracked nuclei (of diameter $8\mu m$) over one hour of live imaging to estimate cell velocities. The resulting velocities were averaged to determine a mean velocity denoted $\overline{v}$. Finally, we use the corresponding tissue density computed from the DAPI embryos (explained in Section 3.1) denoted $\rho$ and compute the number of injected cells per hour in each tissue using the following formula:

$$\text{Number of injected cells per hour} = \rho \times \overline{v} \times w.$$

To obtain the model parameters $\kappa_1$ and $\kappa_2$ we divide by the length of the analyzed injection region, that is $100\mu m$.

#### 3.3 Velocity quantification and sliding

We used live movies of transgenic quail embryos to quantify cell velocity within each tissue (NT and PSM) and inter-tissue sliding. First we changed the reference frame from the laboratory frame to the frame of the last formed somite. This can be achieved by computing the average velocity of cells within the last formed somite and shifting the reference frame to correspond to the average velocity computed. To obtain cell velocities in each of the PSM and of the NT, we first create a reference axis along the antero-posterior axis, specifically going from the last formed somite, denoted $y_{min}$, to the end of the neural tube, denoted $y_{max}$. We denote this distance by $d = y_{max} - y_{min}$. We then detect and track cells (of diameter $8\mu m$) in 5 regions of size $0.1d \times 0.1d \times 10\mu m^3$ equally spaced along the antero-posterior reference axis $[y_{min}, y_{max}]$, over one hour of live imaging. Cell velocities were averaged in each spot, and the velocities at each spot was plotted per embryo (Supplementary Figure 4C-D). Finally, to obtain the sliding, for each embryo, we subtract the average NT velocities obtained from the PSM average velocities and average the resulting sliding to obtain the plot. The same computations were done for the transgenic quail embryo electroporated with p27.

[5] Arthur Michaut. Biomechanics of anteroposterior axis elongation in the chicken embryo. Theses, Université de Strasbourg, September 2018.

[6] Arthur Michaut, Alessandro Mongera, Anupam Gupta, Mattia Serra, Pietro Rigoni, Jong Gwan Lee, Felipe Duarte, Adam R. Hall, L. Mahadevan, Karine Guevorkian, and Olivier Pourquié. Activity-driven extracellular volume expansion drives vertebrate axis elongation. 2022.

[7] Ido Regev, Karine Guevorkian, Anupam Gupta, Olivier Pourquié, and L. Mahadevan. Rectified random cell motility as a mechanism for embryo elongation. Development, 149(6):dev199423, 03 2022.
